# Real-time single-molecule observation of chaperone-assisted protein folding

**DOI:** 10.1101/2022.05.09.491114

**Authors:** Nicholas Marzano, Bishnu Paudel, Antoine van Oijen, Heath Ecroyd

**Affiliations:** Molecular Horizons and School of Chemistry and Molecular Bioscience, University of Wollongong, Wollongong, New South Wales, 2522, Australia; Illawarra Health and Medical Research Institute, Wollongong, New South Wales, 2522, Australia

**Author notes:** **Contact info** Correspondence. Lead contacts (Antoine van Oijen, Heath Ecroyd). equal senior authors (Antoine van Oijen, Heath Ecroyd).

**Keywords:** smFRET, Hsp40, Hsp70, protein folding, single-molecule fluorescence, TIRF microscopy, luciferase, molecular chaperones, protein misfolding

## Abstract

The ability of Hsp70 molecular chaperones to remodel the conformation of their clients is central to their biological function; however, questions remain regarding the precise molecular mechanisms by which Hsp70 machinery interacts with the client and how this contributes towards efficient protein folding. Here, we used Total Internal Reflection Fluorescence (TIRF) microscopy and single-molecule Fluorescence Resonance Energy Transfer (smFRET) to temporally observe the conformational changes that occur to individual firefly luciferase (Fluc) proteins as they are folded by the bacterial Hsp70 system. For the first time, we observed multiple cycles of chaperone binding-and-release to an individual client during refolding and that high rates of chaperone cycling improves refolding yield. Furthermore, we demonstrate that DnaJ remodels misfolded proteins via a conformational selection mechanism whereas DnaK resolves misfolded states via mechanical unfolding. This study illustrates that the temporal observation of chaperone-assisted folding enables the elucidation of key mechanistic details inaccessible using other approaches.

## Introduction

Most proteins fold into specific three-dimensional conformations in order to carry out their unique biological roles (Dobson, 2003). However, during periods of cellular stress or in response to genetic mutation, proteins can become misfolded. This process can contribute to the progression of diseases associated with protein loss-of-function (e.g., Cystic fibrosis) (Fraser-Pitt *et al*., 2015) or the accumulation of protein aggregates, as occurs in Parkinson’s disease, Alzheimer’s disease and cataracts (Dauer *et al*., 2003; Truscott, 2005; Thal *et al*., 2015). Consequently, cells have evolved a complex network of molecular chaperones that act to prevent protein misfolding and maintain protein homeostasis (Taylor *et al*., 2005; Hartl *et al*., 2011; Chiti *et al*., 2017). The Hsp70 family of molecular chaperones represents a central hub of this network, performing a plethora of cellular roles including *de novo* protein folding and refolding, protein disaggregation, protein translocation and the assembly and disassembly of protein complexes (Calloni *et al*., 2012; Goloubinoff, 2017). The mechanisms by which Hsp70 folds client proteins are highly regulated by its co-chaperones Hsp40 and nucleotide-exchange factors (NEFs).

Much of our current understanding regarding the mechanisms by which Hsp70 folds its client proteins has been elucidated through the bacterial system that comprises a single Hsp70 (DnaK), Hsp40 (DnaJ) and NEF (GrpE) (Szabo *et al*., 1994; Wisén *et al*., 2008; Bonomo *et al*., 2010; Luengo *et al*., 2018). The current dogma of Hsp70 function is that protein folding is initiated upon binding of a misfolded or unfolded client by Hsp40, which acts as a ‘holdase’ until the client is delivered to the ATP-bound Hsp70 ‘foldase’. Binding of the Hsp40-client complex to Hsp70 stimulates the rate of hydrolysis of ATP to ADP by more than 1,000 fold (Laufen *et al*., 1999; Russell *et al*., 1999), which results in high-affinity capture of the bound client by Hsp70. Binding of a NEF to ADP-bound Hsp70 accelerates the release of ADP, which results in concomitant rebinding of ATP and returns Hsp70 to the low-affinity ‘open conformation’. Consequently, the client protein dissociates from Hsp70 and can either spontaneously refold to the native state or become misfolded and be subjected to additional rounds of chaperone binding and release.

Owing to the heterogeneous nature of chaperone-client interactions and the conformational state of the client, fundamental questions remain regarding the precise molecular mechanisms by which chaperones remodel the conformation of their clients. For instance, it is unclear whether chaperones stabilize unfolded conformers within a misfolded ensemble (e.g., conformational selection) (Sekhar *et al*., 2018) or whether they utilize ATP hydrolysis to mechanically unfold client proteins (in the case of Hsp70) (Sharma *et al*., 2010; Kellner *et al*., 2014; Dahiya *et al*., 2019; Imamoglu *et al*., 2020). Furthermore, it remains to be determined whether chaperones promote protein folding by introducing structural and entropic constraints on the polypeptide that minimize non-native interactions or smooth folding landscapes. Such mechanistic insights are crucial in our understanding of how chaperones work to fold proteins but are typically difficult to elucidate using conventional ensemble-averaging approaches. This limitation is largely because the rare and transient states that are inherently present during chaperone-assisted folding are masked by billions of asynchronous molecules visiting those states at different times.

Single-molecule approaches have become increasingly popular as a tool to better understand the fundamental mechanisms of chaperone function owing to their capacity to directly observe single molecules and detect kinetic features and states that are normally hidden. Consequently, such approaches have been applied to study how molecular chaperones affect the conformational state of their client proteins and assist in their refolding (Kellner *et al*., 2014; Sekhar *et al*., 2015; Tsuboyama *et al*., 2018; Imamoglu *et al*., 2020). However, while these previous studies have provided valuable insights, most of them did not allow for long-term visualization of the folded state of the same molecule.

In this work we have, for the first time, directly observed the conformational changes that occur to individual client proteins as they are folded by the bacterial Hsp70 system (e.g., DnaK, DnaJ and GrpE) with temporal resolution. To do so, we developed a firefly luciferase (Fluc) construct that can be immobilized to a coverslip surface such that the conformation of individual Fluc molecules can be monitored over time using Total Internal Reflection Fluorescence (TIRF) microscopy and single-molecule Fluorescence Resonance Energy Transfer (smFRET). By analyzing the kinetic data generated from FRET trajectories of individual molecules, we demonstrate that DnaJ remodels misfolded proteins via a conformational selection mechanism whereas DnaK conformationally expands the client via mechanical unfolding. We provide direct evidence that Fluc undergoes repeated cycles of chaperone binding-and-release during refolding and that the rate of chaperone cycling correlates with an improved refolding yield.

## Methods

### Resource availability

#### Lead contact

Further information and requests for resources and reagents should be directed to and will be fulfilled by the Lead Contacts, Heath Ecroyd and Antoine van Oijen.

#### Materials availability

All unique reagents generated in this study are available from the lead contact without restriction.

#### Data and code availability

Single-molecule and other raw data materials are available from the lead contact upon request. All original code has been deposited at Zenodo and is publicly available as of the date of publication at the following DOI:10.5281/zenodo.6392452. Any additional information required to re-analyze the data reported in this paper is available from the lead contact upon request.

### Materials

Recombinant wild-type Fluc (Fluc^WT^) was purchased from Sigma-Aldrich (Missouri, USA). Plasmids encoding GST-tagged Fluc mutants were generously donated by Prof. Till Böcking (UNSW, Australia). An N-terminal SUMO solubility tag and a C-terminal AviTag were added to the Fluc mutant through standard cloning procedures. Plasmids encoding DnaK, DnaJ and GrpE chaperones were generously donated by Prof. Matthias Mayer (University of Heidelberg, Germany).

### Rationale and design of Fluc smFRET construct

To employ Fluc as a sensor of protein folding using smFRET, we used a Fluc construct in which all native cysteines were substituted with alanines (C81A, C391A) and two cysteine residues were introduced for fluorescent labelling with a donor and acceptor fluorophore pair. The introduced cysteines are positioned on the external faces of the N-terminal domain (K141C) and C-terminal domain (K491C) of Fluc and span either side of the interdomain cleft (Figure 1A). Based on the crystal structure of the protein (PDB 1LCI, Conti *et al*., 1996), these cysteines are 34.8 Å apart (measured from C*α* – C*α*), which enables the relative structure and positioning of both Fluc domains to be monitored via smFRET. The modified Fluc construct (i.e., Fluc^ΔCys, K141C, K491C^) is herein denoted as Fluc^IDS^, since the introduced cysteines enable this construct to be used as an interdomain sensor of protein folding. The C-terminal domain of Fluc folds efficiently and independently (does not require chaperones), while the N-terminal domain can often misfold into a state that is rate-limiting for folding (Scholl *et al*., 2014); owing to the different capacities by which these domains fold, this Fluc^IDS^ construct reports on the overall structure of Fluc at the endpoint of folding.

**Figure 1:**
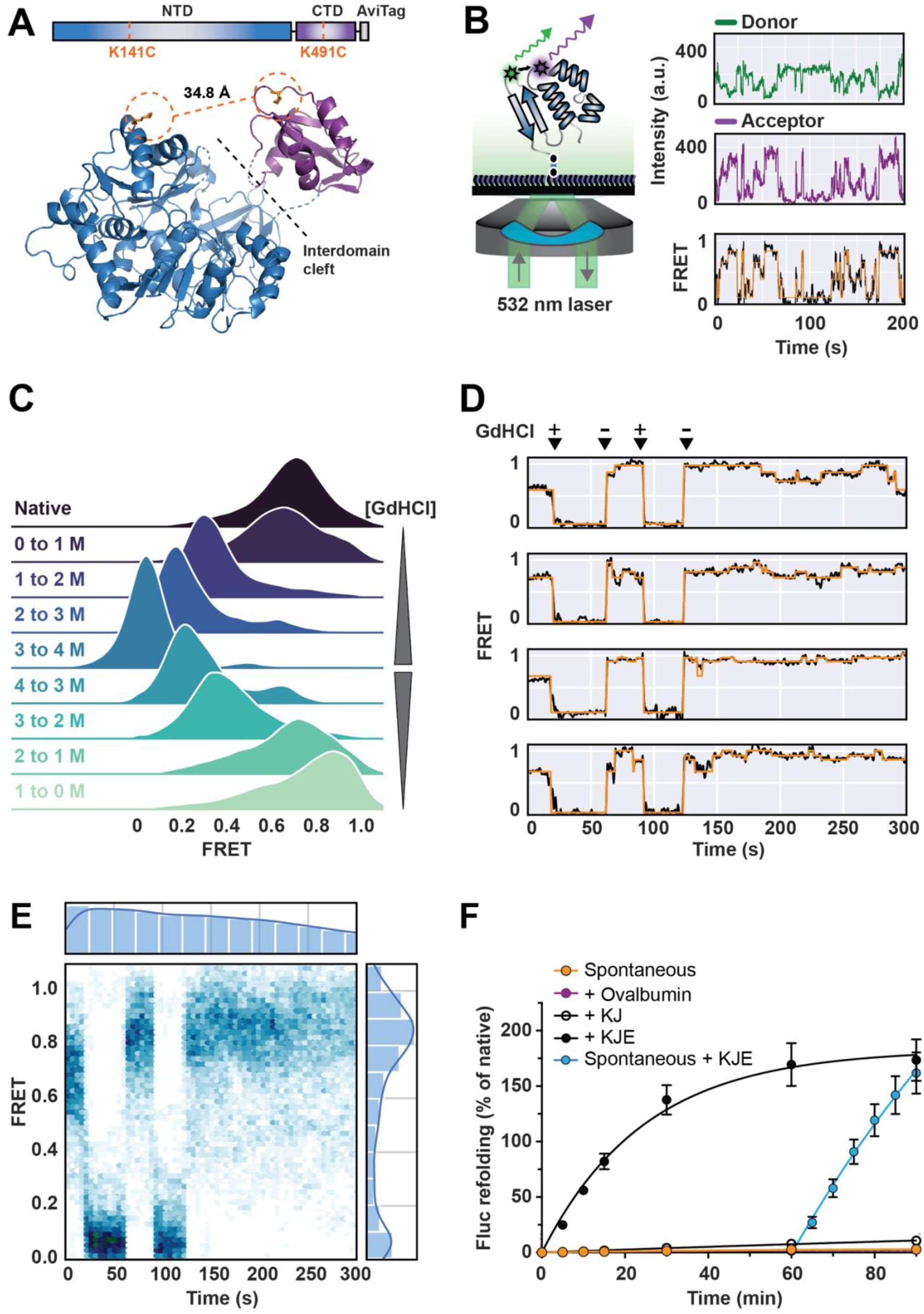
The conformation of individual Fluc^IDS^ molecules can be monitored in real time using smFRET and can be refolded by the bacterial Hsp70 system. **(A)** Schematic of the Fluc^IDS^ FRET-construct employed in this work (*top*) and the structure of Fluc^WT^ (PDB 1LCI, Conti *et al*., 1996) (*bottom*). Residues that were mutated to cysteines for labelling with a FRET fluorophore pair and the C*α* – C*α* distance between labelled residues in the Fluc^IDS^ construct are shown. **(B)** Schematic of the smFRET imaging setup. AF555/AF647-labelled Fluc^IDS^ is immobilized to a neutravidin-functionalized coverslip surface via the biotinylated C-terminal AviTag. Immobilized protein is illuminated by a 532 nm laser that selectively excites the AF555 donor fluorophore and the fluorescence from both AF555 and AF647 (acceptor) fluorophores are measured over time and the FRET efficiency calculated. **(C)** Ridgeline plot of the FRET efficiency of Fluc^IDS^ molecules when unfolded by increasing concentrations of GdHCl (0 – 4 M) and during refolding following dilution out of denaturant. **(D - E)** Immobilized Fluc^IDS^ was incubated in imaging buffer supplemented with or without 4 M GdHCl and the FRET efficiency was measured every 500 ms. **(D)** Representative smFRET traces of individual Fluc^IDS^ molecules. The addition (+) and removal (-) of 4 M GdHCl from the flow cell is indicated above the traces. **(E)** The FRET efficiency over time for all Fluc^IDS^ molecules were collated and plotted as a heatmap (n = 92 molecules). **(F)** Fluc^IDS^ in 5 M GdHCl was diluted 100-fold (10 nM final concentration) into refolding buffer alone (i.e., spontaneous) or buffer supplemented with molecular chaperones or the non-chaperone control protein, ovalbumin (15 µM). Chaperone-assisted refolding was performed with DnaK (3 µM) and DnaJ (1 µM) only (*+ KJ*) or in combination with GrpE (1.5 µM) (*+ KJE*). Rescue of misfolded Fluc^IDS^ was performed by 10-fold dilution of spontaneously refolded Fluc^IDS^ (1 nM final concentration) into refolding buffer containing DnaK (3 µM), DnaJ (1 µM) and GrpE (1.5 µM) (*spontaneous + KJE*). All treatments contained 5 mM ATP. Data shown represent the mean ± standard error of the mean from three independent experiments.

### Expression and purification of recombinant protein

#### Fluc^IDS^

*Escherichia coli* (*E. coli*) BL21(DE3) cells co-transformed with plasmids encoding biotin ligase (BirA) and Fluc^IDS^ (with N-terminal 6x-His-SUMO and C-terminal AviTag motifs) were used to inoculate a starter culture consisting of LB media supplemented with kanamycin (50 µg/mL) and chloramphenicol (10 µg/mL) and grown overnight at 37°C with constant agitation at 180 rpm in an orbital shaker. The starter culture was used to inoculate expression cultures containing LB media supplemented only with kanamycin (50 µg/mL) and the cultures incubated at 37°C until an OD_600_ of ∼ 0.4 was reached. Expression cultures were then further incubated at 18°C until an OD_600_ of ∼ 0.6 - 0.8 was reached. To promote the *in vivo* biotinylation of the AviTagged recombinant protein, the media was supplemented with D-biotin (50 µM final concentration) prepared in 10 mM bicine buffer (pH 8.3) prior to protein induction. Expression of recombinant protein was induced by addition of isopropyl-β-D-1-thiogalactopyranoside (IPTG, 0.1 mM), with the cultures incubated on an orbital shaker at 130 rpm overnight (∼ 20 hr) at 18°C. The cells were then harvested by centrifugation at 5,000 x *g* for 10 min at 4°C and the pellet stored at −20°C until the recombinant protein was extracted.

Recombinant Fluc^IDS^ was extracted from the bacterial pellet via resuspension in 50 mM Tris-HCl (pH 8.0) supplemented with 300 mM NaCl, 5 mM imidazole, 10% (v/v) glycerol (IMAC buffer A) that also contained lysozyme (0.5 mg/mL) and EDTA-free cocktail protease inhibitor. The resuspended pellet was then incubated at 4°C for 20 min and the lysates subjected to probe sonication for 3 min (10 s on/20 s off) at 45% power. The cell debris was then pelleted twice at 24,000 x *g* for 20 min at 4°C and the soluble bacterial lysate passed through a 0.45 µm filter to remove particulates prior to subsequent purification.

The lysate containing recombinant Fluc^IDS^ (with N-terminal 6x-His-SUMO tag) was loaded onto a 5 mL His-Trap Sephadex column pre-equilibrated in IMAC buffer A and bound protein eluted with IMAC buffer A supplemented with 500 mM imidazole (IMAC buffer B). Fractions containing eluted protein were pooled and dialyzed overnight at 4°C in the presence of Ulp1 (4 µg/mg of recombinant protein) against IMAC buffer A that did not contain imidazole (IMAC buffer C). Recombinant Fluc^IDS^ was further purified away from contaminant proteins and the cleaved SUMO tag by passing the dialysate over the same His-Trap Sephadex IMAC column equilibrated in IMAC buffer C and was purified as previously described, however, this time the recombinant protein did not bind to the column and so was collected in flow through fractions. Fractions containing recombinant protein were pooled and dialyzed into 10 mM Tris (pH 9.0) supplemented with 0.5 mM EDTA and 10% (v/v) glycerol (IEX buffer A) for further purification by ion-exchange (IEX) chromatography. Dialyzed protein was loaded onto a MonoQ 5/50 column pre-equilibrated in IEX buffer A. Recombinant Fluc^IDS^ was eluted from the column using a linear salt gradient (0 – 300 mM NaCl) over 20 column volumes. Eluate fractions containing recombinant protein were pooled and dialyzed into storage buffer (50 mM Tris [pH 7.5], 10% (v/v) glycerol), concentrated, snap-frozen in liquid nitrogen and stored at −80°C until required.

#### Molecular chaperones

BL21(DE3) *E. coli* cells transformed with plasmid encoding bacterial chaperones (i.e., DnaK, DnaJ and GrpE) were used to inoculate a starter culture consisting of LB media supplemented with the appropriate antibiotic (50 µg/mL kanamycin for DnaK and DnaJ, 100 µg/mL ampicillin for GrpE) and grown overnight at 37°C. The cultures were used to inoculate expression cultures containing LB media with the appropriate antibiotic and were grown at 37°C to the required OD_600_ (0.8 for DnaK, 1.2 for DnaJ or GrpE). Recombinant protein expression was induced upon the addition of 1 mM IPTG (for DnaK or DnaJ) or 0.2% (w/v) L-(+)-arabinose (for GrpE) and cells harvested following 4 hr of shaking at 30°C. The bacterial pellets were resuspended in protein-specific lysis buffers (DnaK: 20 mM Tris [pH 7.9], 100 mM KCl, 20 mM imidazole; DnaJ: 50 mM Tris [pH 7.9], 500 mM NaCl, 0.1% (v/v) Triton X100, 5 mM MgCl_2_; GrpE: 20 mM Tris [pH 7.9], 100 mM NaCl, 5 mM MgCl_2_) that also contained lysozyme (0.5 mg/mL) and EDTA-free cocktail protease inhibitor. Recombinant protein was extracted, and the lysate clarified, as described above for Fluc^IDS^.

All proteins were purified using IMAC. Briefly, the lysate containing recombinant DnaK was passed over an IMAC column equilibrated in the appropriate lysis buffer and washed in lysis buffer supplemented first with 20 mM imidazole and then with 5 mM ATP and 5 mM MgCl_2_. Bound protein was eluted in lysis buffer supplemented with 250 mM imidazole, with bound fractions pooled and dialyzed against lysis buffer in the presence of Ulp1 (4 µg/mL of recombinant protein) to remove the N-terminal 6x-His-SUMO tag. The dialysate was then passed onto the IMAC column a second time and purified as described above, with fractions containing recombinant protein pooled, dialyzed into storage buffer (40 mM HEPES-KOH [pH 7.6], 100 mM KCl, 5 mM MgCl_2_, 10 mM BME, 10% (v/v) glycerol) and stored at −80°C until use. Recombinant GrpE was purified as described above for DnaK, with the exception that the GrpE lysis buffer was used instead of the DnaK lysis buffer and recombinant GrpE was dialyzed into 40 mM HEPES-KOH (pH 7.6) supplemented with 150 mM NaCl, 10% (v/v) glycerol for storage at −80°C.

DnaJ was purified as described for DnaK using the appropriate lysis buffer with the following modifications. Lysate containing recombinant DnaJ was loaded onto an IMAC column equilibrated in the appropriate lysis buffer and washed in lysis buffer supplemented first with 1.5 M NaCl and then supplemented with 200 mM NaCl. Bound protein was then eluted in lysis buffer supplemented with 250 mM imidazole and 200 mM NaCl, with bound fractions pooled and dialyzed against lysis buffer in the presence of Ulp1 (4 mg total) to remove the N-terminal 6x-His-SUMO tag. The dialysate was then passed onto the IMAC column a second time and purified as described above, with fractions containing recombinant protein pooled, dialyzed into storage buffer (40 mM HEPES-KOH [pH 7.6], 300 mM KCl, 10% (v/v) glycerol) and stored at −80°C until use.

### Refolding assays

The ability of denatured Fluc^IDS^ to spontaneously refold to a native state was assessed by measuring the return of enzymatic activity after denaturation. Denatured Fluc^IDS^ was prepared by incubation of native Fluc^IDS^ in unfolding buffer (50 mM HEPES-KOH [pH 7.5], 50 mM KCl, 5 mM MgCl_2_, 2 mM DTT and 5 M guanidinium hydrochloride [GdHCl]) for 30 min at room temperature. Spontaneous refolding of denatured Fluc^IDS^ was initiated by a 1:100 dilution into refolding buffer (50 mM HEPES-KOH [pH 7.5], 80 mM KCl, 5 mM MgCl_2_, 2 mM DTT and 0.05% (v/v) Tween-20) such that the final concentration of Fluc^IDS^ was 10 nM. Refolding reactions with bacterial chaperones (i.e., DnaK, DnaJ and GrpE, denoted as the KJE system) were left to incubate at 25°C for up to 90 min. Throughout the refolding reactions, aliquots were taken at various times and dispensed into the bottom of a white 96-well CoStar plate (Sigma-Aldrich, USA). The luminescence reaction was initiated following injection of a 10-fold excess of assay buffer (25 mM glycylglycine [pH 7.4], 0.25 mM luciferin, 100 mM KCl, 15 mM MgCl_2_ and 2 mM ATP) into a single well and, 5 s after injection, the luminescence was measured for 10 s using a PolarStar Omega (BMG Labtech, Germany) plate reader. The injection and measurement procedure described above was then performed sequentially for each well to ensure consistency between the measurements. All measurements were performed at 25°C with the gain set to 4,000. Refolding yields were calculated by normalizing to the activity of native (non-denatured) Fluc^IDS^, which was treated as described above with the exception that GdHCl was omitted from the unfolding buffer.

Chaperone-assisted refolding reactions were performed as described above with the exception that denatured Fluc^IDS^ was diluted into refolding buffer containing chaperones and nucleotide. Briefly, refolding assays containing bacterial chaperones were performed with either Fluc^IDS^ or Fluc^WT^ in the presence of 1 µM DnaJ, 3 µM DnaK and 1.5 µM GrpE (when present) supplemented with either 2 mM or 5 mM ATP. To monitor the KJE-assisted refolding of misfolded Fluc^IDS^, following 60 min of unassisted refolding (i.e., spontaneously refolded treatment), an aliquot of Fluc^IDS^ was diluted 10-fold (1 nM final concentration) into refolding buffer containing the KJE chaperone system and the luminescence was measured at various timepoints. All data were fit to a one-phase association curve to determine the plateau (denoted as *Y*_max_) and time required to achieve half *Y*_max_ (denoted as *t*_1/2_) using GraphPad Prism 9 software.

### Total internal reflection fluorescence (TIRF) microscopy

#### Protein labelling

Fluc^IDS^ was fluorescently labelled with an Alexa Fluor 555 (AF555) and Alexa Fluor 647 (AF647) FRET-pair as described previously (Kim *et al*., 2008) with minor modifications. Briefly, Fluc^IDS^ (∼ 2 mg/mL) was incubated in the presence of 5 mM tris(2-carboxyethyl)phosphine (TCEP) and 40% (w/v) ammonium sulfate, and placed on a rotator for 1 hr at 4°C. Proteins were then centrifuged at 20,000 x *g* for 15 min and resuspended in degassed buffer A (100 mM Na_2_PO_4_ [pH 7.4], 200 mM NaCl, 1 mM EDTA and 40% (w/v) ammonium sulfate). The protein was then centrifuged at 20,000 x *g* for 15 min and resuspended in degassed buffer B (100 mM Na_2_PO_4_ [pH 7.4], 200 mM NaCl, 1 mM EDTA). Fluc^IDS^ was then incubated in the presence of a 4-fold and 6-fold excess of pre-mixed AF555 donor and AF647 acceptor fluorophores, respectively, and placed on a rotator overnight at 4°C. Following the coupling reaction, excess dye was removed by gel filtration chromatography using a 7K MWCO Zeba Spin Desalting column (Thermo Fisher Scientific, USA) equilibrated in 50 mM Tris (pH 7.5) supplemented with 20% (v/v) glycerol. The concentration and degree of labelling were calculated by UV absorbance.

#### Microscope setup

Samples were imaged using a custom-built TIRF microcopy system constructed using an inverted optical microscope (Nikon Eclipse TI) that was coupled to an electron-multiplied charge-coupled device (EMCDD) camera (Hamamatsu Photonics K. K., Model C9100-13, Japan). The camera was integrated to operate in an objective-type TIRF setup with diode-pumped solid-state lasers (200 mW Sapphire; Coherent, USA) emitting circularly polarized laser radiation of either 488, 532 or 647-nm continuous wavelength. The laser excitation light was reflected by a dichroic mirror (ZT405/488/532/647; Semrock, USA) and directed through an oil-immersion objective lens (CFI Apochromate TIRF Series 60x objective lens, numerical aperture = 1.49) and onto the sample. Total internal reflection was achieved by directing the incident ray onto the sample at an angle at the critical angle (*θ*_c_) of ∼ 67**°** for a glass/water interface. The evanescent light field generated selectively excites the surface-immobilized fluorophores, with the fluorescence emission passing through the same objective lens and filtered by the same dichroic mirror. The emission was then passed through a 635-nm long pass filter (BLP01-635R; Semrock, USA) and the final fluorescent image projected onto the EMCDD camera. The camera was running in frame transfer mode at 20 Hz, with an electron multiplication gain of 700, operating at − 70°C with a pixel distance of 110 nm (in the sample space).

#### Coverslip preparation and flow cell assembly

Coverslips were functionalized with neutravidin as previously described (Chandradoss *et al*., 2014), with minor modifications. Briefly, 24 x 24 mm glass coverslips were cleaned by alternatively sonicating in 100% ethanol and 5 M KOH for a total of 2 hr before aminosilanization in 2% (v/v) 3-aminopropyl trimethoxysilane (Alfa Aesar, USA) for 15 min. NHS-ester methoxy-polyethylene glycol, molecular weight 5 kDa (mPEG) and biotinylated-mPEG (bPEG; LaysanBio, USA), at a 20:1 (w/w) ratio, was dissolved in 50 mM 4-morpholinepropanesulfonic acid (MOPS, pH 7.4) buffer and sandwiched between two activated coverslips for a minimum of 4 hr for initial passivation in a custom-made humidity chamber. PEGylated coverslips were then rinsed with milli-Q and PEGylated again as described above for overnight (∼ 20 hr) passivation. PEGylated coverslips were rinsed with milli-Q water, dried under nitrogen gas, and stored at −20°C until required. Prior to use, neutravidin (0.2 mg/mL; BioLabs, USA) in milli-Q was incubated on the passivated coverslip for 10 min to bind to the bPEG. Neutravidin functionalized coverslips were then rinsed with milli-Q, dried under nitrogen gas and bonded to a polydimethylsiloxane (PDMS) flow cell for use in single-molecule experiments. Finally, to reduce the non-specific binding of proteins to the coverslip surface, each channel in the microfluidic setup was incubated in the presence of 2% (v/v) Tween-20 for 30 min as previously described (Pan *et al*., 2015) and then washed with imaging buffer.

#### Surface immobilization of labelled proteins and acquisition of smFRET data

For smFRET experiments, labelled Fluc^IDS^ was specifically immobilized to a neutravidin-functionalized and Tween-20-coated coverslip. To do so, labelled protein (∼ 50 pM final concentration) was diluted in imaging buffer (50 mM Tris [pH 7.5], 80 mM KCl, 10 mM MgCl_2_, 200 mM BME and 6 mM 6-hydroxy-2,5,7,8-tetramethylchroman-2-carboxylic acid [Trolox]) and incubated in the flow cell for 5 min. These conditions would typically give rise to ∼ 200 – 300 FRET-competent molecules per 100 x 100 µm. Unbound proteins were then removed from the flow cell by flowing through imaging buffer.

All data were acquired using the TIRF microscope setup previously described following sample illumination using a 532-nm solid state laser with excitation intensity of 2.6 W/cm^2^ and the fluorescence of donor and acceptor fluorophores was measured every 200 ms or 500 ms (for GdHCl-induced folding and unfolding experiments) at multiple fields of view. An oxygen scavenging system (OSS) consisting of 5 mM protocatechuic acid and 50 nM protocatechuate-3,4-dioxygenase was included in all buffers prior to and during image acquisition to minimize photobleaching and fluorophore blinking.

### smFRET data experiments

#### Characterization of the Fluc^IDS^ FRET sensor

To characterize the ability of Fluc^IDS^ to report on changes in conformation via changes in FRET efficiency in real time, single-molecule experiments were performed in which the folded state of Fluc^IDS^ was perturbed with the chemical denaturant GdHCl. To monitor protein unfolding, labelled Fluc^IDS^ incubated in imaging buffer was imaged in the absence of denaturant (i.e., native state) and after 5 min incubation with increasing concentrations of GdHCl (0 – 4 M). The refolding of Fluc^IDS^ denatured in 4 M GdHCl was observed by imaging immediately following the addition of decreasing concentrations of GdHCl (3 – 0 M). To demonstrate the ability of Fluc^IDS^ to report on structural transitions in real time, immobilized Fluc^IDS^ molecules were alternatively incubated in imaging buffer supplemented with or without 4 M GdHCl and the FRET efficiency measured.

#### Monitoring the refolding of Fluc^IDS^ by the bacterial Hsp70 system

To assess the effect of molecular chaperones on the conformation of a client protein, Fluc^IDS^ was incubated in 4 M GdHCl for 5 min and the denaturant removed with imaging buffer to generate a misfolded state. To assess the effect of the DnaJ co-chaperone on the conformation of misfolded Fluc^IDS^, the FRET efficiency of individual molecules were imaged in the presence of increasing concentrations of DnaJ (0 – 10 µM). To assess the effect of DnaK on the conformation of the client, misfolded Fluc^IDS^ was incubated in the presence of DnaJ (0.2 µM), ATP (5 mM) and increasing concentrations of DnaK (0 – 10 µM). As a control, the FRET efficiency of Fluc^IDS^ was determined in the presence of DnaK alone (3 µM) or when supplemented with DnaJ (0.2 µM) and the non-hydrolysable ATP analog, AMP-PNP (5 mM). Finally, the conformation of Fluc^IDS^ during refolding was assessed by incubation of misfolded protein with DnaJ (0.6 µM), DnaK (9 µM) and different concentrations of GrpE (0 – 6 µM). Each KJE-assisted refolding reaction was performed in a different microfluidic channel and smFRET data was acquired for 36 min post-addition of the complete KJE system to misfolded Fluc^IDS^.

### smFRET analysis

Single-molecule time trajectories were analyzed in MATLAB using the MASH-FRET user interface (https://rna-fretools.github.io/MASH-FRET/) (Hadzic *et al*., 2018). The approximate FRET value is measured as the ratio between the acceptor fluorescence intensity (*I_Acceptor_*) and the sum of both donor (*I_Donor_*) and acceptor fluorescence intensities after correcting for crosstalk between donor and acceptor channels. The formula for calculating the FRET efficiency is given by Equation 1, whereby the corrected acceptor intensity (denoted as *CI_Acceptor_*) is equal to *I_Acceptor_ –* (γ * *I_Donor_*) and γ is the crosstalk correction constant. γ is calculated as the ratio of fluorescence measured in the acceptor and donor detection channels following direct excitation of a protein labelled with a single donor fluorophore.

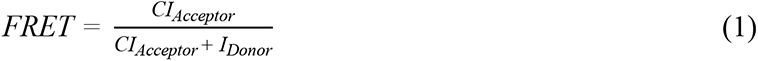

Briefly, donor and acceptor fluorescence channels were aligned following a local weighted mean transformation of images of TetraSpeck fluorescence beads and donor and acceptor fluorescence spots co-localized to identify FRET pairs. Molecules that displayed clear donor and/or acceptor photobleaching events or demonstrated clear anti-correlated changes in donor and acceptor fluorescence intensity were selected for subsequent analysis. The number of photobleaching events observed were used to determine the number of fluorophores present; only molecules in which a single donor and acceptor photobleaching event was observed were used for further analysis. Selected molecules were smoothed using the NL filter coded in MASH-FRET and data truncated to only include FRET values acquired before donor or acceptor photobleaching. FRET efficiency data were exported to the state finding algorithm vbFRET (https://sourceforge.net/projects/vbFRET/) and trajectories fit to a Hidden Markov Model (HMM) to identify discrete FRET states and the transition distributions between them. Default vbFRET settings were employed to fit data to the HMM, with the exception that the *mu* and *beta* hyperparameters were changed to 1.5 and 0.5, respectively, to prevent over-fitting.

Since it is not possible to determine for how long a particular FRET state would have existed if not truncated due to photobleaching, the last measured residence times were deleted from the dataset to extract true residence times. Since data were smoothed during denoising, residence times shorter than that given by Equation 2 were not considered for further analysis. *F* is the imaging frame rate in milliseconds and *N_FA_* is the number of frames that were averaged during trace denoising.

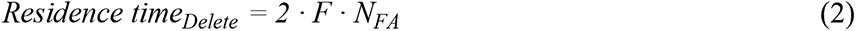

The FRET efficiency of all datapoints for each molecule was collated and is presented as FRET efficiency histograms. To identify the proportion of the different Fluc^IDS^ states within the smFRET data (e.g., Hsp70-bound, native or misfolded), histograms were fit to a mixed gaussian model. Native Fluc^IDS^ was fit to a normally distributed gaussian. Due to the non-linear relationship between dye-pair distances and the FRET efficiency, FRET populations with theoretical dye-pair distances that are significantly deviated from the Förster radius (5.1 nm for AF555/AF647), such as Hsp70-bound and misfolded Fluc^IDS^ populations, need to be fit to skewed gaussian models (Supplementary Figure 1). Native or misfolded states of Fluc^IDS^ were fit to their respective gaussian shapes in the absence of chaperones and the *sigma*, *center*, and *gamma* (for the skewed gaussian model) parameter values were used as a guide when analyzing smFRET data acquired in the presence of chaperones. The area under each curve was determined and normalized to the combined area of all gaussians to determine the proportion of each Fluc^IDS^ state.

To characterize the changes in Fluc^IDS^ conformation that occur during imaging, the frequency of FRET transition distributions was plotted as a function of the FRET state before (*F*_Before_) and after (*F*_After_) each transition as a transition density plot (TDP, Supplementary Figure 2). To investigate transitions of interest, each transition was sorted into different directional classes denoted generally as T_Before-After_, whereby ‘before’ refers to *F*_Before_ and ‘after’ refers to *F*_After_. For simplicity, FRET data were filtered according to whether *F*_Before_ or *F*_After_ was greater than 0.5 (high) or less than 0.5 (low), unless otherwise indicated, and thus four different transition classes are possible: T_high-high_, T_high-low_, T_low-low_ and T_low-high_. Finally, the occurrence of each transition class was normalized to the total number of transitions measured and the corresponding mean residence time (defined as the time that a molecule resides at *F*_Before_ prior to transition to *F*_After_) for each transition class was calculated (Supplementary Figure 2). The residence times were statistically analyzed using a one-way or two-way analysis of variance (ANOVA) with either a Šídák or Tukey’s multiple comparisons post-hoc test, with p ≤ 0.05 determined to be statistically significant. All data analysis and presentation were performed using custom-written scripts on Python software or using GraphPad Prism 9 (GraphPad Software Inc; San Diego, USA).

## Results and discussion

### The conformation of Fluc^IDS^ can be monitored in real time using smFRET and TIRF microscopy

Fluc was chosen for this work as it possesses several characteristics that make it an ideal client protein for the study of chaperone function; Fluc readily misfolds following dilution from chemical denaturant, has a limited ability to spontaneously fold or refold (it requires chaperones to do so) and the attainment of its native state can be readily assessed via a highly sensitive bioluminescence enzymatic assay (Levy *et al*., 1995; Nillegoda *et al*., 2015; Hristozova *et al*., 2016). We first characterized the ability of the Fluc^IDS^ construct to report on conformational changes in real time (Figure 1A). To do so, Fluc^IDS^ was immobilized to a neutravidin-functionalized coverslip and imaged in the presence of different concentrations of denaturant using TIRF microscopy and microfluidics (Figure 1B). Native Fluc^IDS^ resulted in a FRET distribution centered at ∼ 0.7 (Figure 1C), which decreased with increasing concentrations of denaturant such that, when in 4 M GdHCl, the FRET distribution was centered at 0.1. Such a result is consistent with the transition of Fluc^IDS^ to a conformationally expanded random-coil structure that has been observed previously in other single-molecule unfolding experiments (Schuler *et al*., 2002; Kuzmenkina *et al*., 2005; Tezuka-Kawakami *et al*., 2006; Rosenkranz *et al*., 2011). Progressive dilution of denaturant resulted in an increase in FRET efficiency; however, complete removal of Fluc^IDS^ out of denaturant resulted in Fluc^IDS^ having a FRET efficiency that was higher than that of the native state (∼ 0.9 vs ∼ 0.7, respectively). This high-FRET state corresponds to a previously observed misfolded Fluc conformation, which is thought to occur as a result of the formation of non-native contacts between residues in the N-terminal domain and/or residues in the C-terminal domain (Mashaghi *et al*., 2014; Scholl *et al*., 2014; Imamoglu *et al*., 2020).

As a proof of principle for the ability of Fluc^IDS^ to act as a real-time sensor of protein folding, individual immobilized Fluc^IDS^ molecules were alternatively incubated in the absence or presence of chemical denaturant. The FRET intensity originating from individual Fluc^IDS^ species was determined and data from all molecules were collated into an averaged heatmap (Figure 1D-E). Native Fluc^IDS^ (measured between 0 – 20 s) exhibited a FRET efficiency of ∼ 0.7, which rapidly dropped to ∼ 0.1 for all Fluc^IDS^ molecules measured upon addition of GdHCl. Removal of GdHCl caused the FRET efficiency to rapidly increase to the high-FRET misfolded state (∼ 0.9). Similar transitions to low-FRET and high-FRET states were observed following a second round of injection into and removal of GdHCl from the flow cell. Together, these data highlight that smFRET is an ideal tool to monitor the real-time conformational changes of Fluc^IDS^, and that Fluc^IDS^ can be used as a sensor of protein folding.

Ensemble-based assays have established that the bacterial Hsp70 system (i.e., DnaK, DnaJ and GrpE) can efficiently refold chemically denatured Fluc (Szabo *et al*., 1994; Wisén *et al*., 2008; Bonomo *et al*., 2010; Luengo *et al*., 2018). As such, we sought to employ this model chaperone system and characterize its ability to refold Fluc^IDS^ in preparation for smFRET experiments. When chemically denatured Fluc^IDS^ was refolded in the absence of chaperones (i.e., spontaneous refolding) or a non-chaperone control protein, ovalbumin, minimal refolding was observed (Figure 1F). However, Fluc^IDS^ was rapidly and efficiently refolded by the KJE system. Crucially, chaperone-assisted refolding was observed to be efficient only when the NEF, GrpE, was present during the reaction. Data from this work and others (Sharma *et al*., 2010; Scholl *et al*., 2014; Imamoglu *et al*., 2020) have shown that the misfolded (high-FRET) Fluc conformation is stable, enzymatically inactive and persists for many hours. Interestingly, however, the addition of the bacterial Hsp70 system to spontaneously misfolded Fluc^IDS^ resulted in rapid and efficient refolding. Collectively, these results demonstrate that the bacterial Hsp70 system efficiently resolves misfolded states of Fluc^IDS^ and refold it to a functional state, and that GrpE is essential for efficient refolding.

Notably, the amount of spontaneous refolding of Fluc^IDS^ (3.6% of the activity of the native [non-denatured] state) was much lower than previously reported for the spontaneous refolding of Fluc^WT^ (e.g., 58.2% by Imamoglu *et al*., 2020). To determine if the low spontaneous refolding yield of Fluc^IDS^ is due to the introduced mutations in this isoform of the protein (i.e., C-terminal AviTag and cysteine substitutions), the refolding experiments were repeated using both Fluc^IDS^ and Fluc^WT^ (Supplementary Figure 3A). Refolding was performed with lower concentrations of ATP as this slows refolding rates and thus augments differences in the refolding kinetics between these two proteins. Indeed, under these conditions significantly more Fluc^WT^ was able to spontaneously refold to the native state (29.6%) compared to Fluc^IDS^ (2.9%). As expected, dilution of Fluc from denaturant in the presence of the bacterial Hsp70 system resulted in the efficient refolding of both Fluc^WT^ and Fluc^IDS^, although the rate of refolding was ∼ 4-fold faster for Fluc^WT^ (*t_1/2_* = 15.9 min) compared to Fluc^IDS^ (*t_1/2_* = 66.0 min).

### DnaJ binds to and stabilizes expanded conformations of Fluc^IDS^ via a conformational selection mechanism

We next sought to use smFRET to investigate how the bacterial Hsp40 homolog, DnaJ, affects the folded state of Fluc^IDS^. When Fluc^IDS^ was allowed to refold in the absence of chaperones, it adopted the previously described high-FRET (∼ 0.8) misfolded conformation (Figure 2A-B). However, when incubated with progressively higher concentrations of DnaJ, the FRET distributions became broader, and a second low-FRET peak centered at ∼ 0.2 – 0.3 was observed. The reduced FRET efficiency of Fluc^IDS^ molecules is consistent with the DnaJ-mediated conformational expansion of client proteins observed previously (Kellner *et al*., 2014).

**Figure 2:**
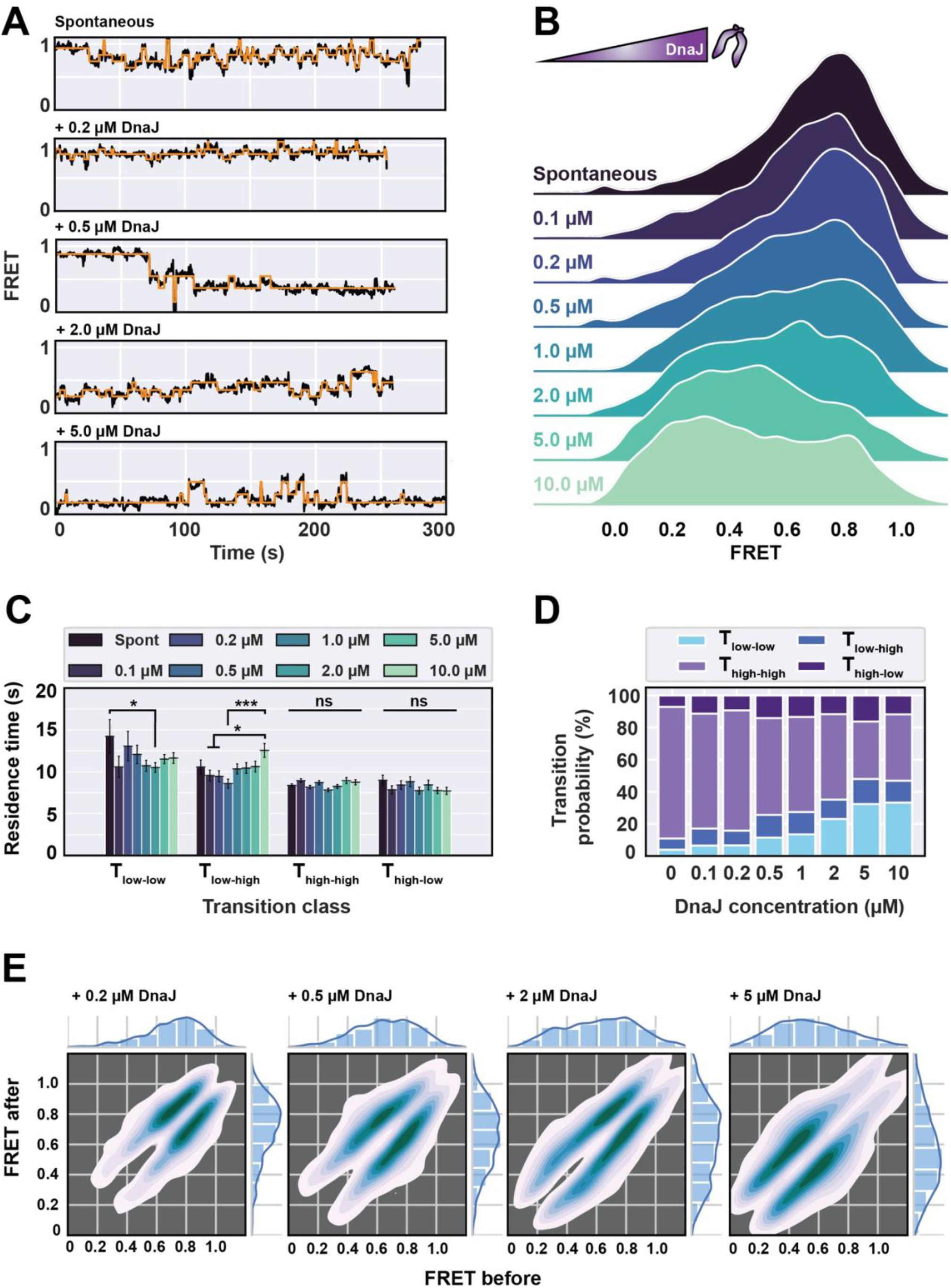
DnaJ stabilizes conformationally expanded states of Fluc^IDS^. Fluc^IDS^ was diluted from denaturant in the presence of increasing concentrations of DnaJ (0 - 10 µM) and the FRET efficiency was measured. **(A)** Representative smFRET traces of individual Fluc^IDS^ molecules when incubated in the absence (i.e., spontaneous) or presence of increasing concentrations of DnaJ (0.2 – 5 µM). Data are fit with an HMM (shown in *orange*). **(B)** Ridgeline plot of the FRET efficiency distributions of misfolded Fluc^IDS^ alone or when incubated with the indicated concentration of DnaJ. **(C)** Bar plots showing the residence time of different transition classes. Data shown is the mean ± standard error of the mean. A two-way ANOVA statistical analysis with Tukey’s multiple-comparisons post-hoc test was performed to determine statistically significant differences in residence times between treatment groups within each transition class. * and *** indicates statistical significance with p ≤ 0.05 and 0.001, respectively. ns or the absence of markers indicates no significant difference (p > 0.05). **(D)** The transition probability of each transition class. **(E)** TDPs of Fluc^IDS^ transitions when incubated in the absence or presence of the indicated concentration of DnaJ. Data for all panels were derived and collated from the HMM fits of at least 268 individual Fluc^IDS^ molecules per treatment.

Kinetically, there are two possible explanations for the increased occupancy of Fluc^IDS^ at low-FRET states when incubated in the presence of DnaJ; (i) low-FRET states are longer-lived than high-FRET states; or (ii) transitions that result in the accumulation of low-FRET states of Fluc^IDS^ are favored. Kinetic analysis of individual FRET trajectories demonstrates that the residence times of Fluc^IDS^ molecules are relatively unaffected with increasing concentrations of DnaJ, with the exception that there is an increase in residence time for low-FRET to high-FRET transitions (i.e., T_low-high_) at the highest concentration of DnaJ tested (10 µM) (Figure 2C). Conversely, higher concentrations of DnaJ significantly increased the occurrence of transitions between low-FRET states of Fluc^IDS^ (i.e., T_low-low_) (Figure 2D); thus, Fluc^IDS^ remains conformationally dynamic but is constrained within an ensemble of non-native and conformationally expanded states when bound to DnaJ. Such conformational dynamicity and/or destabilization of Hsp40-bound client proteins have been observed previously, whereby HDX experiments demonstrated that amide hydrogens exchange more rapidly upon binding of Fluc by DnaJ (Rodriguez *et al*., 2008). Furthermore, a recent study found that the client binding sites on many Hsp40 isoforms spans across the C-terminal subdomain of both Hsp40 homodimers (Jiang *et al*., 2019). Spatial separation of the client across both Hsp40 homodimers results in the stabilization of expanded non-native conformations of the client, prevents spontaneous collapse and may dissolve secondary and tertiary structural elements of the bound client (Jiang *et al*., 2019), results consistent with this work.

It remains to be established whether DnaJ selectively binds to a subset of already expanded conformations of Fluc^IDS^ (e.g., a conformational selection model), or if binding by DnaJ remodels the conformation of the client protein to expanded states (e.g., an induced fit model) (Goloubinoff *et al*., 2007; Rodriguez *et al*., 2008; Sharma *et al*., 2010; Mayer, 2013; Clerico *et al*., 2015; Sekhar *et al*., 2018). Kinetic analysis of the FRET efficiency traces from individual Fluc^IDS^ species demonstrate that most FRET transitions occur between high-FRET values (i.e., 0.6 – 1.0) in the presence of low concentrations of DnaJ (≤ 0.5 µM), while the transition density shifts to lower FRET states (i.e., 0.3 – 0.6) with higher concentrations of DnaJ (i.e., ≥ 2 µM) (Figure 2E). Notably, the transition distributions are symmetrically mirrored and close to the TDP diagonal-axis at all concentrations of DnaJ tested, which indicates that the observed transitions occur reversibly and are small in amplitude (Figure 2E, Supplementary Figure 3B). Notably, the occurrence of large Fluc^IDS^ conformational changes (> 0.5) does not change with increasing concentrations of DnaJ (Supplementary Figure 3B), which suggests that DnaJ binding does not typically induce mechanical unfolding of the client. Since the residence times of T_high-low_ transitions do not significantly change with DnaJ concentration and the TDPs do not demonstrate transition density away from the diagonal-axis, our data support a model in which DnaJ scans non-native states and binds to exposed hydrophobic regions typically exposed in already expanded client conformations (Rüdiger *et al*., 2001). Although Hsp40 chaperones typically form complexes with their client proteins at a 1:1 ratio (Terada *et al*., 2010; Kellner *et al*., 2014; Jiang *et al*., 2019), the observation that T_low-high_ residence times are longer in the presence of high concentrations of DnaJ suggests that multiple chaperones can bind to a single misfolded client (since dissociation of individual species is concentration-independent). Thus, these data demonstrate that the binding of multiple DnaJ proteins to a single client result in the stabilization of unfolded conformers and partitions the client towards a conformationally expanded ensemble.

### DnaK entropically pulls clients to resolve misfolded states

Next, we sought to investigate how DnaJ co-operates with DnaK to fold Fluc^IDS^. To assess the sole effect of DnaK on the conformation of Fluc^IDS^, increasing concentrations of DnaK were incubated with a concentration of DnaJ (0.2 µM) that alone does not significantly affect the conformation of Fluc^IDS^ but is sufficient to promote DnaK loading onto the client (Supplementary Figure 3C). Incubation of Fluc^IDS^ with DnaJ only, or when supplemented with a low concentration of DnaK (0.2 µM), did not significantly affect the FRET traces or distributions of Fluc^IDS^ (Figure 3A-B). Incubation of misfolded Fluc^IDS^ with increasing concentrations of DnaK resulted in the re-distribution of high-FRET populations (> 0.5) towards an ultra-low FRET state (defined as < 0.2) such that, at the highest concentration of DnaK, the ultra-low FRET peak constituted > 70% of the total FRET species. FRET intensity traces from Fluc^IDS^ molecules incubated with higher concentrations of DnaK (≥ 2 µM) demonstrate that these ultra-low FRET states are stable and can persist for minutes (Figure 3A). As expected, the formation of this ultra-low FRET state is dependent on the presence of DnaJ and ATP hydrolysis and thus constitutes a DnaK-bound and conformationally expanded form of Fluc^IDS^ (Supplementary Figure 3C). It is interesting, however, that DnaJ can promote binding of DnaK to Fluc^IDS^ at a DnaJ concentration that by itself is not sufficient to significantly remodel the conformational landscape of Fluc^IDS^. Such a result suggests that remodeling of the client by DnaJ is not strictly necessary during refolding and that its predominant role is to stimulate ATP hydrolysis and client capture by DnaK; the conformational expansion of the client at high DnaJ concentrations is thus likely a mechanism by which it can prevent the well described misfolding and subsequent aggregation of Fluc (Szabo *et al*., 1994; Perales-Calvo *et al*., 2010). Importantly, the ultra-low FRET conformationally-expanded conformation of Fluc^IDS^ constitutes a necessary intermediate for chaperone-mediated on-pathway folding, since the absence of ATP from refolding assays inhibits productive Fluc^IDS^ refolding. Collectively, these results demonstrate that the DnaK-bound Fluc^IDS^ molecules are more conformationally expanded compared to DnaJ-bound molecules and contributes to a growing body of literature that conformational expansion of the client is a generic mechanism of Hsp70 function (Kellner *et al*., 2014; Dahiya *et al*., 2019; Imamoglu *et al*., 2020).

**Figure 3:**
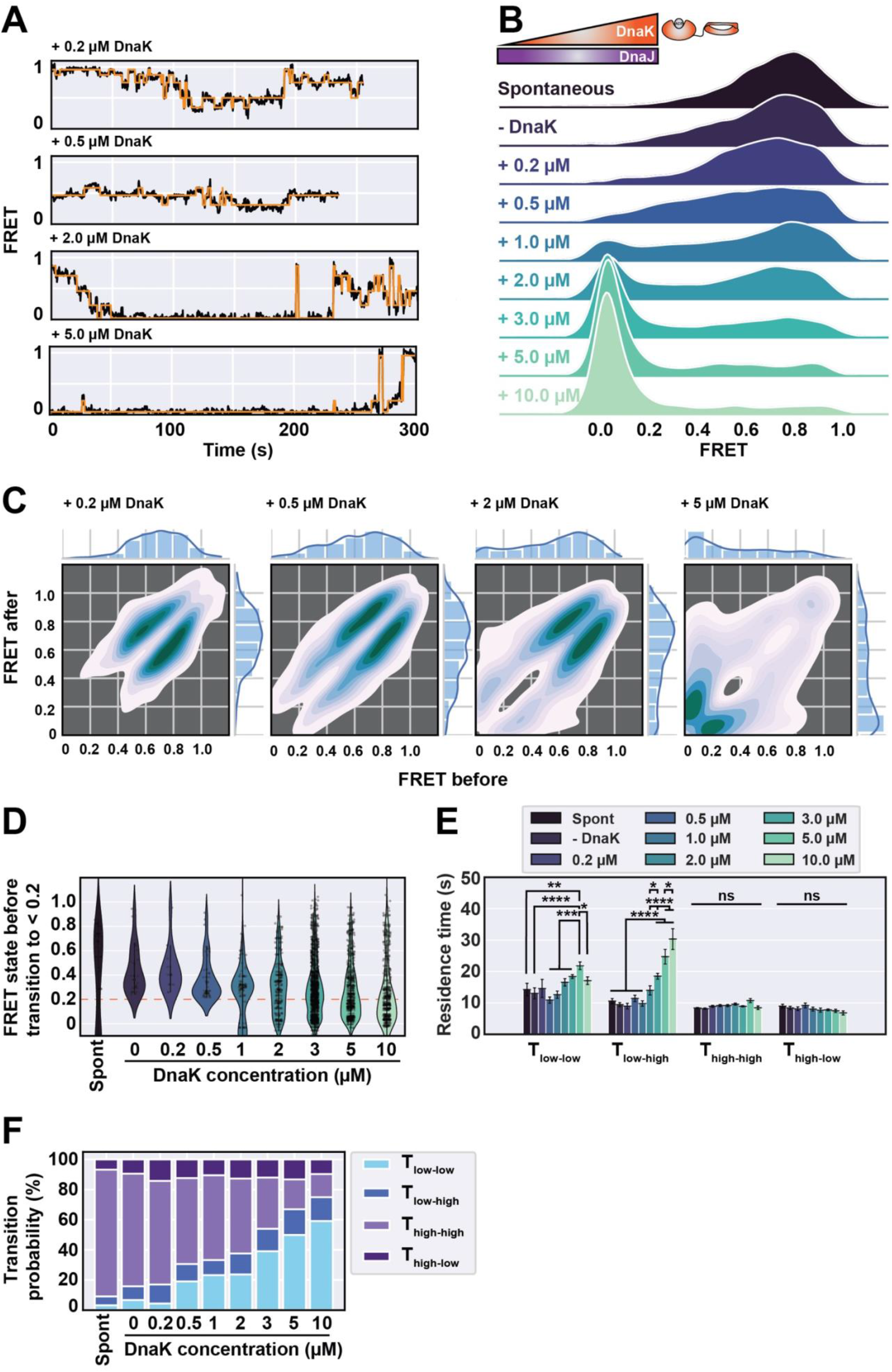
DnaK resolves misfolded states of Fluc^IDS^ by conformational expansion. Fluc^IDS^ was diluted from denaturant in the absence (i.e., spontaneous) or presence of DnaJ (0.2 µM) and increasing concentrations of DnaK (0 – 10 µM) and the FRET efficiency measured. ATP (5 mM) was present for all conditions tested. **(A)** Representative smFRET traces of individual Fluc^IDS^ molecules when incubated in the presence of DnaJ and the indicated concentration of DnaK. Data are fit with an HMM (shown in *orange*). **(B)** Ridgeline plot of the FRET efficiency distributions of misfolded Fluc^IDS^ or when incubated with DnaJ and the indicated concentration of DnaK. **(C)** TDPs showing Fluc^IDS^ transitions when incubated in the presence of DnaJ and the indicated concentration of DnaK. **(D)** Violin plot of the FRET efficiencies of Fluc^IDS^ molecules immediately preceding a transition (i.e., *F*_Before_) to an ultra-low FRET state (< 0.2). **(E)** Bar plots showing the residence time of different transition classes. A two-way ANOVA statistical analysis with Tukey’s multiple-comparisons post-hoc test was performed to determine statistically significant differences in residence times between treatment groups within each transition class. *, **, *** and **** indicates statistical significance with p ≤ 0.05, 0.01, 0.001 and 0.0001, respectively. ns or the absence of markers indicates no significant difference (p > 0.05). **(F)** The transition probability of each transition class. Data for all panels were derived and collated from the HMM fits of at least 155 individual Fluc^IDS^ molecules per treatment.

We next interrogated the kinetic data to reveal the mechanism by which DnaK remodels the conformation of clients. Notably, the TDPs demonstrate that there is a clear difference in the mechanism by which DnaK binds to misfolded protein relative to DnaJ. When Fluc^IDS^ was allowed to refold from denaturant in the absence or presence of low concentrations of DnaK (≤ 0.5 µM), most transitions occur between high-FRET values (i.e., 0.6 – 1.0) and lay close to the diagonal-axis (Figure 3C). Incubation of Fluc^IDS^ with higher concentrations of DnaK (i.e., ≥ 2 µM) results in increased transition density at ultra-low FRET states (e.g., ∼ 0 – 0.4). However, and in contrast to those results observed with DnaJ, the TDPs at high concentrations of DnaK exhibited strong transition densities away from the diagonal-axis which implies the presence of significant Fluc^IDS^ conformational changes. Indeed, high concentrations of DnaK, but not DnaJ, increase the occurrence and proportion of molecules that experience transitions larger in amplitude than 0.5 FRET (Supplementary Figure 3B). There has been some conjecture regarding the mechanism conjecture regarding the mechanism by which Hsp70 resolves misfolded states of proteins (Sharma *et al*., 2010; Kellner *et al*., 2014; Sekhar *et al*., 2018; Dahiya *et al*., 2019; Imamoglu *et al*., 2020); thus, we decided to investigate this by quantifying the FRET state of Fluc^IDS^ immediately prior to a transition to a DnaK-bound state (i.e., < 0.2 FRET) (**Error! Reference source not found.**D). Interestingly, under conditions in which the formation of the DnaK-bound state is favored (≥ 2 µM of DnaK), most transitions originate from ultra-low FRET states. However, a significant proportion of transitions originate from high-FRET states (> 0.5), indicating that DnaK can forcibly unfold compact conformations of Fluc^IDS^.

It is interesting that the formation of DnaK-bound Fluc^IDS^ complexes often originated following a single transition from a high-FRET, compact and misfolded conformation of Fluc^IDS^. Although the conversion of misfolded Fluc^IDS^ to DnaK-bound states occurred following single transition events, which might suggest the binding of an individual DnaK molecule, it remains possible that multiple DnaK species associate with Fluc^IDS^ since only one structural coordinate can be monitored using conventional two-color smFRET. Indeed, considering that the dissociation of a single DnaK species from Fluc^IDS^ is concentration-independent (Kellner *et al*., 2014), the kinetic data supports a model by which multiple DnaK species are bound to Fluc^IDS^ since the apparent dissociation of DnaK from Fluc^IDS^ is significantly slower with increasing concentrations of DnaK (as indicated by longer residence times for T_low-high_ transitions) (Figure 3E). Notably, when the residence times of those transitions that originate from or to DnaK-bound states (i.e., < 0.2 FRET) were determined, a significant 2-fold reduction in residence time was observed for T_high-low_ transitions (Supplementary Figure 3D). Such data demonstrate that the loading of multiple DnaK molecules onto a single client is promoted by increased association rates at high chaperone concentrations, consistent with reports that have shown that up to 12 DnaK species can bind to a single client protein (Kellner *et al*., 2014; Imamoglu *et al*., 2020). Moreover, higher concentrations of DnaK further increase the abundance of transitions constrained at low FRET efficiencies (i.e., T_low-low_ transitions) while the converse was true for T_high-high_ transitions (Figure 3F). Consequently, once bound by DnaK, Fluc^IDS^ is unlikely to transition from a low-FRET state (< 0.5) to a high-FRET state (> 0.5) (Supplementary Figure 3E) but remains conformationally dynamic while constrained in the expanded ensemble. Collectively, these kinetic analyses support an entropic pulling mechanism, whereby the rapid association and subsequent repulsion of multiple DnaK species bound to a single client result in its mechanical unfolding, partitioning the client towards conformationally expanded states.

### Multiple cycles of DnaK binding-and-release are essential for productive folding and is exquisitely regulated by GrpE concentration

Finally, the conformation of Fluc^IDS^ as it is folded by the complete KJE system was assessed. As expected, incubation of misfolded Fluc^IDS^ with DnaJ and DnaK in the absence of GrpE resulted in a shift to the ultra-low FRET state characteristic of DnaK-bound Fluc^IDS^ (Figure 4A-B, additional traces in Supplementary Figure 4A). Consistent with the role of GrpE to dissociate ADP and promote client release from DnaK (Langer *et al*., 1992; Szabo *et al*., 1994; Packschies *et al*., 1997; Brehmer *et al*., 2001), the addition of increasing concentrations of GrpE resulted in the DnaK-bound state of Fluc^IDS^ becoming less abundant with Fluc^IDS^ instead occupying FRET distributions typical of native (∼ 0.6) or misfolded (∼ 0.8) states. To quantify the effect of GrpE concentration on Fluc^IDS^ refolding, the relative proportion of DnaK-bound, native and misfolded Fluc^IDS^ populations was determined by fitting the histogram data with a multiple gaussian model (Figure 4C, Supplementary Figure 5A). As expected, the proportion of DnaK-bound Fluc^IDS^ species was observed to decrease exponentially with increasing concentrations of GrpE; however, misfolded Fluc^IDS^ species became significantly more abundant. Such a result is consistent with the destabilization of DnaK-Fluc^IDS^ complexes by GrpE, as evidenced by the significantly reduced residence times and transition frequencies of DnaK-bound states (i.e., T_low-low_ and T_low-high_) compared to in the absence of GrpE (Supplementary Figure 5B-C). Importantly, the proportion of native Fluc^IDS^ molecules increased until a maximum of 45% was reached at intermediate GrpE concentrations (i.e., 2 µM), but decreased at higher concentrations (Figure 4C).

**Figure 4:**
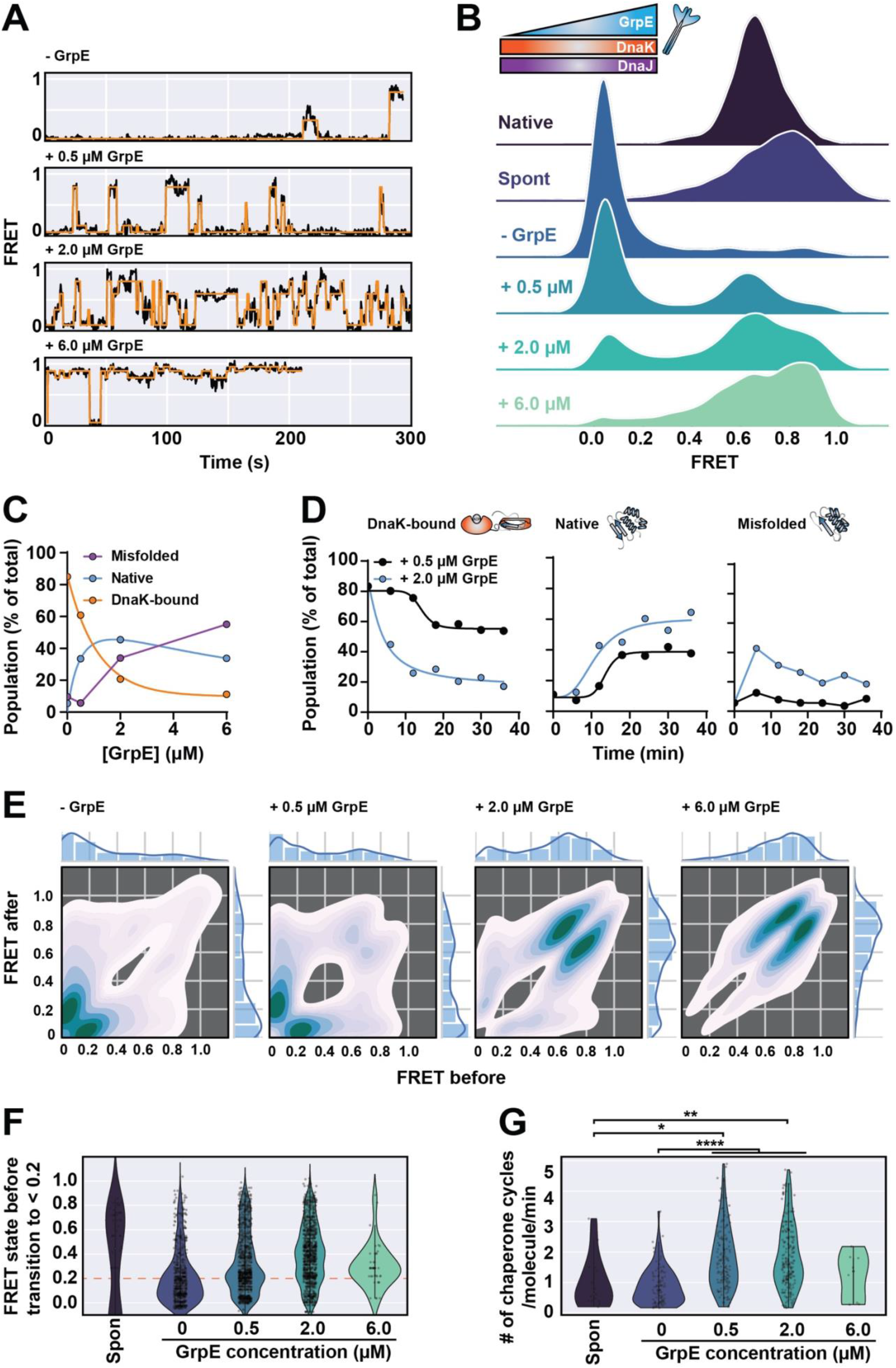
GrpE promotes efficient Fluc^IDS^ refolding by enabling multiple cycles of DnaK binding and release. Fluc^IDS^ was incubated in the presence of DnaJ (0.6 µM), DnaK (9 µM) and ATP (5 mM) supplemented with increasing concentrations of GrpE (0 – 6 µM). FRET efficiency data was measured for 36 min following the addition of GrpE. **(A)** Representative smFRET traces of individual Fluc^IDS^ molecules when incubated in the presence of chaperones. Data are fit with an HMM (shown in *orange*). **(B)** Ridgeline plot of the FRET efficiency distributions of native or misfolded Fluc^IDS^, or when incubated with DnaJ, DnaK and the indicated concentration of GrpE. **(C)** The proportion of Fluc^IDS^ states (i.e., DnaK-bound, native or misfolded) as a function of GrpE concentration. **(D)** The proportion of Fluc^IDS^ states (i.e., DnaK-bound, native or misfolded) over time when incubated in the presence of either 0.5 µM or 2 µM GrpE. **(E)** TDPs showing Fluc^IDS^ transitions when incubated in the presence of DnaJ, DnaK and the indicated concentration of GrpE. **(F)** Violin plots of the FRET efficiencies of Fluc^IDS^ molecules immediately preceding a transition (i.e., *F*_Before_) to an ultra-low FRET state (< 0.2). **(G)** Violin plots showing the distribution of chaperone binding-and-release events to individual Fluc^IDS^ molecules per min in the presence of different concentrations of GrpE. A binding event was classified if *F*_Before_ and *F*_After_ was greater than and less than 0.2 FRET, respectively, and vice-versa for release events. A one-way ANOVA statistical analysis with Tukey’s multiple-comparisons post-hoc test was performed to determine statistically significant differences in the rate of chaperone binding-and-release events between treatment groups. When statistical tests were performed, *, **, *** and **** indicates statistical significance with p ≤ 0.05, 0.01, 0.001 and 0.0001, respectively. ns or the absence of markers indicates no significant difference (p > 0.05). Data for all panels were derived and collated from the HMM fits of at least 205 individual Fluc^IDS^ molecules per treatment.

Such a result suggests that during KJE-mediated refolding there is a compromise between preventing the accumulation of misfolded species and effectively refolding the client to its native state. To interrogate this further, the change in Fluc^IDS^ when incubated with low or intermediate concentrations of GrpE (≤ 2 µM) during refolding was determined (Figure 4D). Low concentrations of GrpE (0.5 µM) delays the opportunity for the client to fold upon chaperone release (as indicated by a high proportion of DnaK-bound states) but reduces the proportion of misfolded species (due to higher DnaK-Fluc^IDS^ affinity): refolding of Fluc^IDS^ to the native state occurs over time but is slow and inefficient, with only 38% of Fluc^IDS^ species occupying native-like FRET distributions. Conversely, intermediate concentrations of GrpE (2 µM) results in the rapid dissociation of DnaK-bound Fluc^IDS^ species, which allows the client an early opportunity to fold correctly. Notably, a high proportion of misfolded protein is generated immediately following initial DnaK release, however, over time the misfolded population is partitioned towards native Fluc^IDS^ states that become the predominant species at later time points.

The current dogma of chaperone-mediated folding suggests that, should the released client remain misfolded, it can undergo additional rounds of chaperone-mediated binding-and-release (Sharma *et al*., 2010; Saibil, 2013; Koldewey *et al*., 2017; Rosenzweig *et al*., 2019; Balchin *et al*., 2020). In this work and for the first time, multiple cycles of chaperone binding-and-release of a single client protein were observed in real time during chaperone-assisted refolding. Many Fluc^IDS^ molecules exhibited repeated transitions to and from ultra-low FRET states when incubated with lower concentrations of GrpE (≤ 2 µM) (> 70%, Figure 4A, additional traces in Supplementary Figure 4B-C, Supplementary Figure 5D), indicative of multiple cycles of DnaK binding-and-release. Accordingly, the TDP analysis demonstrated significant transition density away from the diagonal-axis under these experimental conditions (Figure 4E). Furthermore, a significant proportion of transitions to DnaK-bound states (i.e., < 0.2) were observed to originate from high-FRET misfolded Fluc^IDS^ conformations (> 0.5) during refolding (Figure 4F); such a result supports the TDP analyses and further demonstrates that DnaK can bind to and unfold a variety of misfolded Fluc^IDS^ structures during productive folding. In contrast, very few Fluc^IDS^ molecules experienced transitions to ultra-low FRET states when refolded in the presence of high concentrations of GrpE (6 µM) (< 10%, Figure 4A, Supplementary Figure 5D, additional traces in Supplementary Figure 4D), with the transition density focused in high-FRET regimes along the diagonal-axis of the TDP (Figure 4E).

To ascertain whether multiple chaperone cycles actively promote efficient protein folding, the rate of DnaK binding-and-release to individual client proteins was calculated (Figure 4G). Indeed, under conditions in which Fluc^IDS^ refolding is promoted and misfolding is minimized (i.e., ≤ 2 µM GrpE), the rate of chaperone binding-and-release events per molecule was observed to be significantly higher compared to when refolding is inefficient (i.e., in the absence of GrpE). The requirement for multiple chaperone binding-and-release events for efficient Fluc^IDS^ refolding is further demonstrated by experiments in which only a single binding-and-release event per molecule is possible. To do so, a binding cycle was induced upon incubation of Fluc^IDS^ with high concentrations of DnaJ and DnaK to form DnaK-Fluc^IDS^ complexes and a single round of chaperone release was prompted through introduction into the flow cell a solution containing only GrpE and ATP. After two controlled cycles of chaperone binding-and-release, a native-like FRET distribution was not observed following the GrpE-mediated release of Fluc^IDS^ by DnaK, with Fluc^IDS^ predominantly remaining in the high-FRET misfolded state (Supplementary Figure 5E). These results illustrate that multiple cycles of chaperone binding-and-release are essential for productive Fluc^IDS^ refolding.

## Conclusions

This work sought to address the outstanding question of how the Hsp70 chaperone system promote client refolding. To do so we developed a Fluc construct in which the folded state can be monitored with temporal resolution using a combination of TIRF microscopy and smFRET. For the first time, we have observed the conformation of an individual client protein in real time as it passes through the entire Hsp70 functional cycle and extracted key kinetic and structural details that are typically inaccessible using conventional approaches.

The mechanism by which molecular chaperones interact with clients and affect their conformation is considered central to chaperone function. We show that, unlike DnaK, the binding of multiple DnaJ species to a single client does not result in entropic pulling, but instead prevents the collapse of Fluc^IDS^ to misfolded compact conformations. What then could be the difference in fold-stabilization between these two chaperones? One hypothesis could be that binding of multiple DnaJ species to a single client stabilizes the unfolded conformers; however, since Hsp40 chaperones typically bind to their clients via many weak, low-affinity and multivalent interactions, any instances of steric clashing or kinetic competition between DnaJ molecules for binding sites could accelerate the dissociation of DnaJ species partially bound to Fluc^IDS^. Conversely, DnaK is a non-equilibrium machine that utilizes ATP-hydrolysis to bind clients, which results in the formation of the ADP-bound form of DnaK that is characterized by high affinity for the client (De Los Rios *et al*., 2014). Since dissociation of DnaK is dependent on ADP release, which is rate-limiting in the absence of a NEF, steric clashes between bound DnaK species inherently leads to repulsion of the two chaperones and results in the mechanical unfolding of the client. As such, the relative affinities of DnaJ and DnaK for binding to the client appear to dictate the mechanism by which they affect its conformational landscape; lower affinities favor a conformational selection mechanism while high affinities favor an entropic pulling model. The findings of this study, as well as others (Sharma *et al*., 2010; Imamoglu *et al*., 2020), show that the entropic pulling of client proteins by DnaK helps to resolve folding-incompetent, misfolded structures and that chaperones can conformationally remodel the client for subsequent refolding attempts.

It is traditionally thought that release of the client protein from Hsp70 provides an opportunity for the client to spontaneously refold to the native state. Indeed, recent work has suggested that the role of Hsp70 is to resolve misfolded Fluc states and unfold it (i.e., ‘unfoldase’ activity), following which the client can spontaneously refold to the native state upon release by Hsp70 (i.e., similar to denaturant-induced refolding) (Sharma *et al*., 2010). However, the results of this work suggest that the ‘unfoldase’ mechanism of Hsp70 chaperone function may not fully encompass its role in protein folding and that Hsp70 possesses ‘foldase’ characteristics. As evidence for this, the rate of chaperone-assisted refolding of Fluc^WT^ was ∼ 4-fold faster than that observed for Fluc^IDS^; however, the spontaneous refolding efficiency of Fluc^WT^ was ∼ 10-fold higher than for Fluc^IDS^. This suggests that the chaperone-assisted refolding mechanism extends beyond simple release of an unfolded client protein for spontaneous refolding and that there is some other aspect of chaperone function that accelerates and assists productive folding.

There is a growing body of work that suggests that Hsp70 can direct the folding of proteins in a manner that is distinct from a simple model of binding-and-release (Lee *et al*., 2015; Sekhar *et al*., 2015; Dahiya *et al*., 2019; Lu *et al*., 2021). Kinetic data obtained in this work demonstrated that there was a much lower probability that Fluc^IDS^ would return to high-FRET compacted states once conformationally expanded by Hsp70, indicative of some ‘holdase’-like function. Notably, it has been suggested that the binding of Hsp70 to multiple sites on the client resolves different misfolded structures and may enable the client to occupy heterogeneous and ‘fuzzy’ ensemble structures at regions distinct from the binding site; such structures can subsequently act as folding nuclei that allow the client to explore numerous folding pathways upon chaperone release (Sharma *et al*., 2010; Lee *et al*., 2015; Sekhar *et al*., 2015; Koldewey *et al*., 2017; Dahiya *et al*., 2019). Furthermore, molecular dynamics simulations suggests that Hsp70 may remain transiently associated with the released client and prevent non-native contacts to accelerate productive refolding (Lu *et al*., 2021). While more work is required to validate these hypotheses in the context of Fluc^IDS^ folding, it is clear that Hsp70 can actively remodel the folding landscape of its clients by partitioning them towards native structures and away from non-productive folding pathways.

The kinetic data provides convincing evidence that the refolding efficiency is exquisitely dependent on GrpE concentration and is dictated by several key parameters: i) the relative affinity of DnaK to misfolded Fluc^IDS^ and, ii) the number and rate of DnaK binding-and-release cycles per molecule. The work presented here demonstrates that even low concentrations of GrpE promotes many rapid cycles of DnaK binding-and-release to the client. However, under such conditions the relative affinity of DnaK to Fluc^IDS^ is high (due to slower nucleotide-exchange rates), which results in the accumulation of DnaK onto misfolded Fluc^IDS^ and reduces opportunities for spontaneous refolding. Conversely, high concentrations of GrpE significantly reduce the amount of productive DnaK binding events and thus the number and rate of chaperone binding-and-release events. Thus, refolding is most effective in the presence of intermediate concentrations of GrpE, i.e., when the rate and occurrence of DnaK binding-and-release is high while the relative affinity of DnaK to Fluc^IDS^ remains low.

Collectively, the results described in this work provide a detailed understanding of the mechanism by which the bacterial Hsp70 chaperone system refolds client proteins. Capture of the client by DnaK occurs upon DnaJ-assisted ATP hydrolysis (generating the ADP-bound form of DnaK), which actively drives additional conformational expansion and global unfolding of the bound client. At this stage the client remains conformationally dynamic and may form native contacts in regions distinct from the binding site to prepare for refolding upon DnaK release. Dissociation of ADP from DnaK, which can occur spontaneously but is accelerated by a NEF, results in the concomitant rebinding of ATP and dissociation of the client protein from DnaK. Upon release, DnaK can partition the client protein towards folding pathways en-route to the native state or the client can spontaneously misfold. The misfolded protein can then be subject to additional rounds of binding-and-release by chaperones until the native state is acquired. The results of this work further demonstrate that the rate at which chaperone binding-and-release events occur to a single client is a critical factor in efficient refolding.

## Acknowledgments

We would like to thank Prof. Till Böcking for designing and providing the initial Fluc^IDS^ smFRET construct. We would also like to thank Prof. Matthias Mayer for providing expression plasmids for the bacterial chaperones DnaK, DnaJ and GrpE. We would also like to acknowledge Dezerae Cox for her useful advice when writing the smFRET analysis code. Finally, we thank the staff in Molecular Horizons and the Illawarra Health and Medical Research Institute for their technical and administrative support. This work was funded by the Australian Research Council (DP220103466). AMvO also acknowledges funding from the National Health and Medical Research Council (APP1197069).

## Author contributions

NRM, BPP, HE and AMvO formulated the experimental approach. NRM performed all experiments, curated, and analyzed data, constructed figures, and wrote the initial manuscript. NRM, BPP, HE and AMvO edited the manuscript and approved submission of the final manuscript.

## Declaration of interests

The authors declare no competing interests.

## Notes

### Competing Interest Statement

The authors have declared no competing interest.

## References

Balchin, D., Hayer-Hartl, M. and Hartl, F. U. (2020) ‘Recent advances in understanding catalysis of protein folding by molecular chaperones’, FEBS Letters, 594(17), pp. 2770–2781. doi: 10.1002/1873-3468.13844.

Bonomo, J. et al. (2010) ‘Comparing the functional properties of the Hsp70 chaperones, DnaK and BiP.’, Biophysical Chemistry, 149(1–2), pp. 58–66. doi: 10.1016/j.bpc.2010.04.001.

Brehmer, D. et al. (2001) ‘Tuning of chaperone activity of Hsp70 proteins by modulation of nucleotide exchange’, Nature Structural Biology, 8(5), pp. 427–432. doi: 10.1038/87588.

Calloni, G. et al. (2012) ‘DnaK functions as a central hub in the E. coli chaperone network’, Cell Reports, 1(3), pp. 251–264. doi: 10.1016/J.CELREP.2011.12.007.

Chandradoss, S. D. et al. (2014) ‘Surface passivation for single-molecule protein studies’, Journal of Visualized Experiments, 86(e50549), pp. 1–8. doi: 10.3791/50549.

Chiti, F. and Dobson, C. M. (2017) ‘Protein misfolding, amyloid formation, and human disease: a summary of progress over the last decade’, Annual Review of Biochemistry, 86, pp. 27–68. doi: 10.1146/annurev-biochem.

Clerico, E. M. et al. (2015) ‘How Hsp70 molecular machines interact with their substrates to mediate diverse physiological functions’, Journal of Molecular Biology, 427(7), pp. 1575–1588. doi: 10.1016/j.jmb.2015.02.004.

Conti, E., Franks, N. P. and Brick, P. (1996) ‘Crystal structure of firefly luciferase throws light on a superfamily of adenylate-forming enzymes’, Structure, 4(3), pp. 287–298. doi: 10.1016/S0969-2126(96)00033-0.

Dahiya, V. et al. (2019) ‘Coordinated conformational processing of the tumor suppressor protein p53 by the Hsp70 and Hsp90 chaperone machineries’, Molecular Cell, 74(4), pp. 816–830. doi: 10.1016/j.molcel.2019.03.026.

Dauer, W. and Przedborski, S. (2003) ‘Parkinson’s disease’, Neuron, 39(6), pp. 889–909. doi: 10.1016/S0896-6273(03)00568-3.

Dobson, C. M. (2003) ‘Protein folding and misfolding’, Nature, 426(6968), pp. 884–890. doi: 10.1038/nature02261.

Fraser-Pitt, D. and O’Neil, D. (2015) ‘Cystic fibrosis - a multiorgan protein misfolding disease.’, Future Science, 1(2), p. FSO57. doi: 10.4155/fso.15.57.

Goloubinoff, P. (2017) ‘Editorial: the HSP70 molecular chaperone machines’, Frontiers in Molecular Biosciences, 4(1), pp. 1–4. doi: 10.3389/fmolb.2017.00001.

Goloubinoff, P. and Rios, P. D. L. (2007) ‘The mechanism of Hsp70 chaperones: (entropic) pulling the models together’, Trends in Biochemical Sciences, 32(8), pp. 372–380. doi: 10.1016/J.TIBS.2007.06.008.

Hadzic, M. C. A. S. et al. (2018) ‘Reliable state identification and state transition detection in fluorescence intensity-based single-molecule förster resonance energy-transfer data’, Journal of Physical Chemistry B, 122(23), pp. 6134–6147. doi: 10.1021/acs.jpcb.7b12483.

Hartl, F. U., Bracher, A. and Hayer-Hartl, M. (2011) ‘Molecular chaperones in protein folding and proteostasis’, Nature, 475(7356), pp. 324–332. doi: 10.1038/nature10317.

Hristozova, N., Tompa, P. and Kovacs, D. (2016) ‘A novel method for assessing the chaperone activity of proteins’, PLoS ONE, 11(8), p. e0161970. doi: 10.1371/journal.pone.0161970.

Imamoglu, R. et al. (2020) ‘Bacterial Hsp70 resolves misfolded states and accelerates productive folding of a multi-domain protein’, Nature Communications, 11(1), p. 365. doi: 10.1038/s41467-019-14245-4.

Jiang, Y., Rossi, P. and Kalodimos, C. G. (2019) ‘Structural basis for client recognition and activity of Hsp40 chaperones’, Science, 365(6459), pp. 1313–1319. doi: 10.1126/science.aax1280.

Kellner, R. et al. (2014) ‘Single-molecule spectroscopy reveals chaperone-mediated expansion of substrate protein’, Proceedings of the National Academy of Sciences of the United States of America, 111(37), pp. 13355–13360. doi: 10.1073/pnas.1407086111.

Kim, Y. et al. (2008) ‘Efficient site-specific labeling of proteins via cysteines’, Bioconjugate Chemistry, 19(3), pp. 786–791. doi: 10.1021/bc7002499.

Koldewey, P., Horowitz, S. and Bardwell, J. C. A. (2017) ‘Chaperone-client interactions: Non-specificity engenders multifunctionality’, Journal of Biological Chemistry, 292(29), pp. 12010–12017. doi: 10.1074/jbc.R117.796862.

Kuzmenkina, E. V, Heyes, C. D. and Nienhaus, G. U. (2005) ‘Single-molecule forster resonance energy transfer study of protein dynamics under denaturing conditions’, Proceedings of the National Academy of Sciences of the United States of America, 102(43), pp. 15471–15476. doi: 10.1073/pnas.0507728102.

Langer, T. et al. (1992) ‘Successive action of DnaK, DnaJ and GroEL along the pathway of chaperone-mediated protein folding’, Nature, 356(6371), pp. 683–689. doi: 10.1038/356683a0.

Laufen, T. et al. (1999) ‘Mechanism of regulation of Hsp70 chaperones by DnaJ cochaperones’, Proceedings of the National Academy of Sciences of the United States of America, 96(10), pp. 5452– 5457. doi: 10.1073/pnas.96.10.5452.

Lee, J. H. et al. (2015) ‘Heterogeneous binding of the SH3 client protein to the DnaK molecular chaperone’, Proceedings of the National Academy of Sciences of the United States of America, 112(31), pp. 4206–4215. doi: 10.1073/pnas.1505173112.

Levy, E. J. et al. (1995) ‘Conserved ATPase and luciferase refolding activities between bacteria and yeast Hsp70 chaperones and modulators’, FEBS Letters, 368(3), pp. 435–440. doi: 10.1016/0014-5793(95)00704-D.

Liberek, K. et al. (1991) ‘Escherichia coli DnaJ and GrpE heat shock proteins jointly stimulate ATPase activity of DnaK.’, Proceedings of the National Academy of Sciences of the United States of America, 88(7), p. 2874.

De Los Rios, P. and Barducci, A. (2014) ‘Hsp70 chaperones are non-equilibrium machines that achieve ultra-affinity by energy consumption’, eLife, 3. doi: 10.7554/eLife.02218.

Lu, J. et al. (2021) ‘Energy landscape remodeling mechanism of Hsp70-chaperone-accelerated protein folding’, Biophysical Journal, 120(10), pp. 1971–1983. doi: 10.1016/J.BPJ.2021.03.013.

Luengo, T. M. et al. (2018) ‘Hsp90 breaks the deadlock of the Hsp70 chaperone system’, Molecular Cell, 70(3), pp. 545–552.e9. doi: 10.1016/j.molcel.2018.03.028.

Mashaghi, A., Mashaghi, S. and Tans, S. J. (2014) ‘Misfolding of luciferase at the single-molecule level’, Angewandte Chemie, 53(39), pp. 10390–10393. doi: 10.1002/anie.201405566.

Mayer, M. P. (2013) ‘Hsp70 chaperone dynamics and molecular mechanism’, Trends in Biochemical Sciences, 38(10), pp. 507–514. doi: 10.1016/J.TIBS.2013.08.001.

Nillegoda, N. B. et al. (2015) ‘Crucial HSP70 co-chaperone complex unlocks metazoan protein disaggregation’, Nature, 524(7564), pp. 247–251. doi: 10.1038/nature14884.

Packschies, L. et al. (1997) ‘GrpE accelerates nucleotide exchange of the molecular chaperone DnaK with an associative displacement mechanism’, Biochemistry, 36(12), pp. 3417–3422. doi: 10.1021/BI962835L.

Pan, H. et al. (2015) ‘A simple procedure to improve the surface passivation for single molecule fluorescence studies’, Physical Biology, 12(4), p. 045006. doi: 10.1088/1478-3975/12/4/045006.

Perales-Calvo, J., Muga, A. and Moro, F. (2010) ‘Role of DnaJ G/F-rich domain in conformational recognition and binding of protein substrates.’, Journal of Biological Chemistry, 285(44), pp. 34231– 34239. doi: 10.1074/jbc.M110.144642.

Rampelt, H. et al. (2012) ‘Metazoan Hsp70 machines use Hsp110 to power protein disaggregation’, EMBO Journal, 31(21), pp. 4221–4235. doi: 10.1038/emboj.2012.264.

Rodriguez, F. et al. (2008) ‘Molecular basis for regulation of the heat shock transcription factor σ32 by the DnaK and DnaJ chaperones’, Molecular Cell, 32(3), pp. 347–358. doi: 10.1016/J.MOLCEL.2008.09.016.

Rosenkranz, T. et al. (2011) ‘Native and Unfolded States of Phosphoglycerate Kinase Studied by Single-Molecule FRET’, ChemPhysChem, 12(3), pp. 704–710. doi: 10.1002/cphc.201000701.

Rosenzweig, R. et al. (2019) ‘The Hsp70 chaperone network’, Nature Reviews Molecular Cell Biology, 20(11), pp. 665–680. doi: 10.1038/s41580-019-0133-3.

Rüdiger, S., Schneider-Mergener, J. and Bukau, B. (2001) ‘Its substrate specificity characterizes the DnaJ co-chaperone as a scanning factor for the DnaK chaperone’, EMBO Journal, 20(5), pp. 1042– 1050. doi: 10.1093/emboj/20.5.1042.

Russell, R. et al. (1999) ‘DnaJ dramatically stimulates ATP hydrolysis by DnaK: insight into targeting of Hsp70 proteins to polypeptide substrates’, Biochemistry, 38(13), pp. 4165–4176. doi: 10.1021/bi9824036.

Saibil, H. (2013) ‘Chaperone machines for protein folding, unfolding and disaggregation’, Nature Reviews Molecular Cell Biology, 14(10), pp. 630–642. doi: 10.1038/nrm3658.

Scholl, Z. N., Yang, W. and Marszalek, P. E. (2014) ‘Chaperones rescue luciferase folding by separating its domains’, Journal of Biological Chemistry, 289(41), pp. 28607–28618. doi: 10.1074/jbc.M114.582049.

Schuler, B., Lipman, E. A. and Eaton, W. A. (2002) ‘Probing the free-energy surface for protein folding with single-molecule fluorescence spectroscopy’, Nature, 419(6908), pp. 743–747. doi: 10.1038/nature01060.

Sekhar, A. et al. (2015) ‘Mapping the conformation of a client protein through the Hsp70 functional cycle’, Proceedings of the National Academy of Sciences of the United States of America, 112(33), pp. 10395–10400. doi: 10.1073/pnas.1508504112.

Sekhar, A. et al. (2018) ‘Conserved conformational selection mechanism of Hsp70 chaperone-substrate interactions’, eLife, 7, p. e32764. doi: 10.7554/eLife.32764.

Sharma, S. K. et al. (2010) ‘The kinetic parameters and energy cost of the Hsp70 chaperone as a polypeptide unfoldase’, Nature Chemical Biology, 6(12), pp. 914–920. doi: 10.1038/nchembio.455.

Szabo, A. et al. (1994) ‘The ATP hydrolysis-dependent reaction cycle of the Escherichia coli Hsp70 system DnaK, DnaJ, and GrpE’, Proceedings of the National Academy of Sciences, 91(22), pp. 10345– 10349. doi: 10.1073/pnas.91.22.10345.

Taylor, R. P. and Benjamin, I. J. (2005) ‘Small heat shock proteins: a new classification scheme in mammals’, Journal of Molecular and Cellular Cardiology, 38(3), pp. 433–444. doi: 10.1016/J.YJMCC.2004.12.014.

Terada, K. and Oike, Y. (2010) ‘Multiple molecules of Hsc70 and a dimer of DjA1 independently bind to an unfolded protein’, Applied Microbiology and Biotechnology, 285(22), pp. 16789–16797. doi: 10.1074/jbc.M110.101501.

Tezuka-Kawakami, T. et al. (2006) ‘Urea-induced unfolding of the immunity protein Im9 monitored by spFRET’, Biophysical Journal, 91(5), pp. L42–44. doi: 10.1529/biophysj.106.088344.

Thal, D. R. and Fändrich, M. (2015) ‘Protein aggregation in Alzheimer’s disease: Aβ and τ and their potential roles in the pathogenesis of AD’, Acta Neuropathologica, 129(2), pp. 163–165. doi: 10.1007/s00401-015-1387-2.

Truscott, R. J. W. (2005) ‘Age-related nuclear cataract-oxidation is the key’, Experimental Eye Research, 80(5), pp. 709–725. doi: 10.1016/j.exer.2004.12.007.

Tsuboyama, K., Tadakuma, H. and Tomari, Y. (2018) ‘Conformational activation of argonaute by distinct yet coordinated actions of the Hsp70 and Hsp90 chaperone systems’, Molecular Cell, 70(4), pp. 722–729.e4. doi: 10.1016/j.molcel.2018.04.010.

Wisén, S. and Gestwicki, J. E. (2008) ‘Identification of small molecules that modify the protein folding activity of heat shock protein 70’, Analytical Biochemistry, 374(2), pp. 371–377. doi: 10.1016/J.AB.2007.12.009.

