## Supplementary information for "Real-time single-molecule observation of chaperone-assisted protein folding"

### Author notes

* Lead contacts; # equal senior authors

### Contact info

### Supplementary information

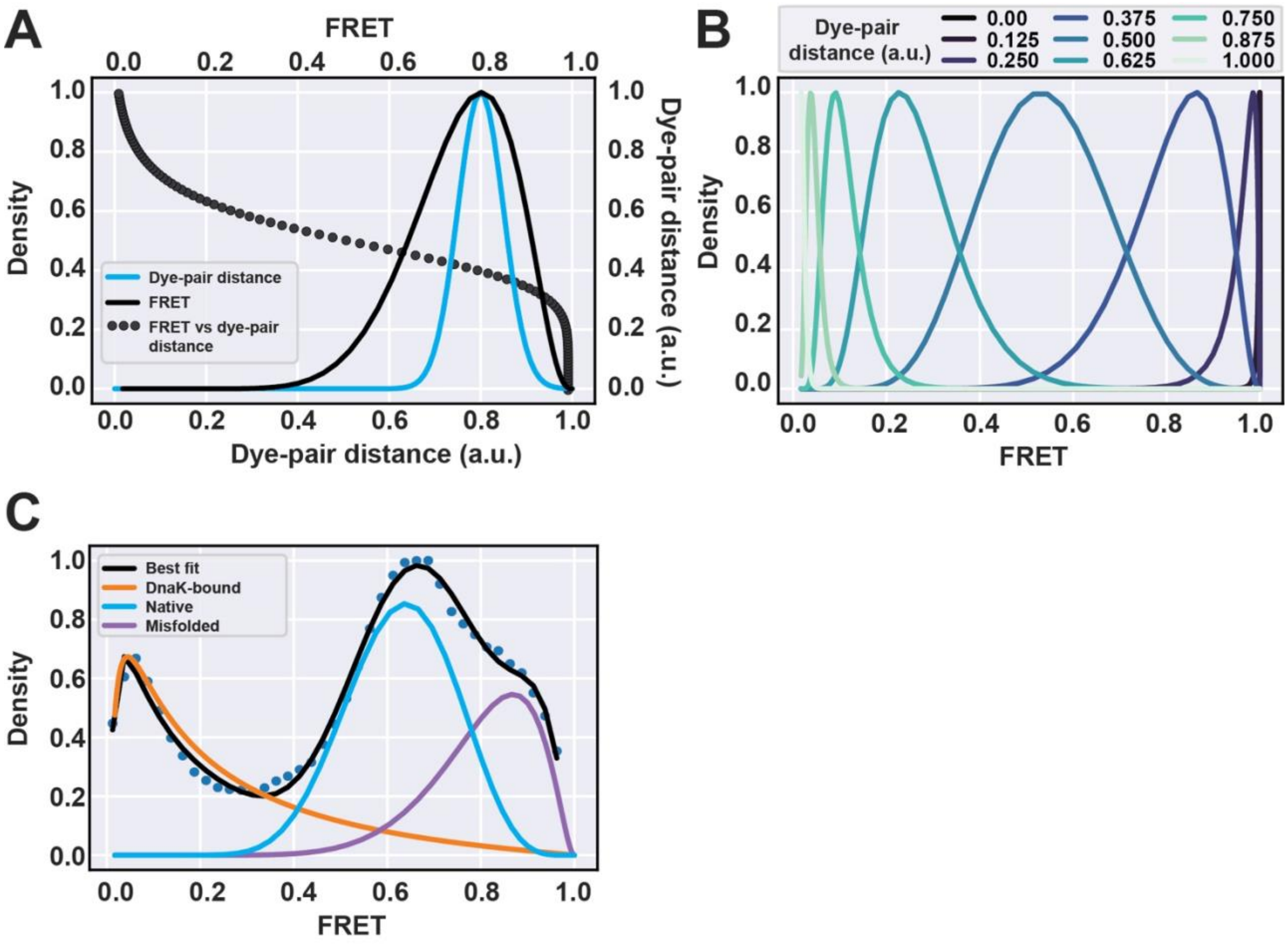


**Supplementary Figure 1: GrpE Non-linear relationship between dye-pair distance and FRET efficiency results in skewed FRET distributions that accurately describe experimental data.** A distribution of dye-pair distances (0 – 1 a.u.) was defined and the theoretical FRET efficiency for each distance was calculated using the FRET equation (Förster radius = 0.51). **(A)** The distribution of dye-pair distances were transformed into a normally distributed Gaussian model and plotted against the dye-pair distance (*blue*, bottom and left axis) or theoretical FRET efficiency (*black*, top and left axis). Each Gaussian was centred at 0.8 FRET with the *sigma*

value set to 0.05. The relationship between the dye-pair distance and the theoretical FRET efficiency (*dotted line*, top and right axis) is also shown. **(B)** Theoretical FRET efficiency distributions when the mean dye-pair distance (*mu*) is centred at different distances. Greater absolute differences of *mu* from the Förster radius results in larger skews and narrower FRET distributions. **(C)** Fits of the experimental smFRET data using the model employed in (B).

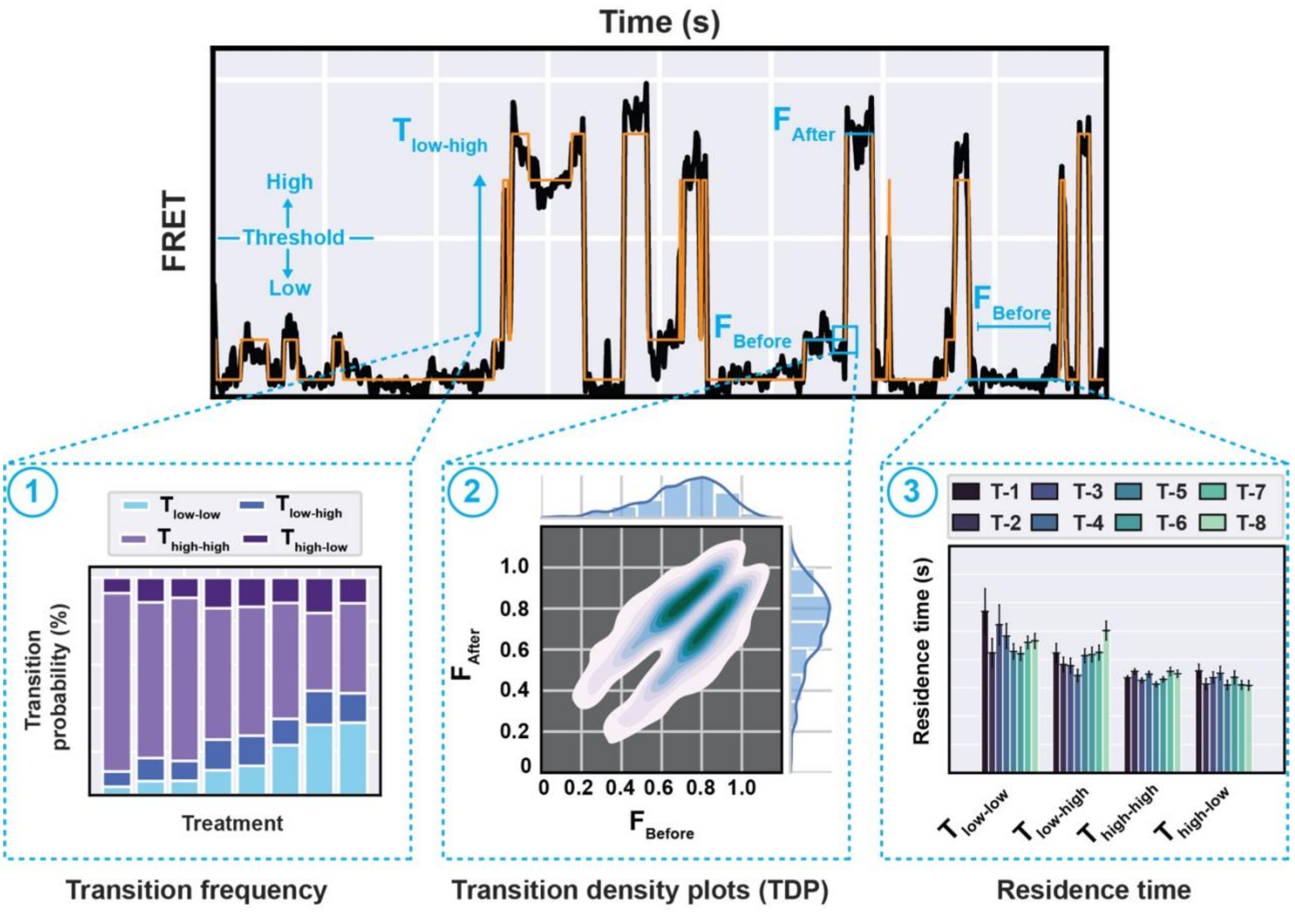


**Supplementary Figure 2: Schematic of the kinetic data acquired from Hidden Markov Model (HMM) fits to the smFRET efficiency data.** An example FRET efficiency trace of a single $\text{Fluc}^{\text{IDS}}$ protein that has been fit with an HMM (*orange*) is shown. For each transition, the FRET state immediately before (defined as $F_{\text{Before}}$) and after (defined as $F_{\text{After}}$) each transition is determined. **(1)** To categorize $\text{Fluc}^{\text{IDS}}$ transitions into distinct classes, the FRET data were filtered according to whether $F_{\text{Before}}$ or $F_{\text{After}}$ was greater than (high) or less than (low) a defined threshold. Typically, the FRET threshold was set to 0.5 unless otherwise indicated. As such, four different transition directions are possible: $T_{\text{high-high}}$, $T_{\text{high-low}}$, $T_{\text{low-low}}$ and $T_{\text{low-high}}$. An example of a $T_{\text{low-high}}$ transition is shown in the schematic. The occurrence of each transition class was normalized to the total number of transitions measured and presented as a cumulative bar plot. **(2)** The frequency of transition distributions can also be plotted as a function of $F_{\text{Before}}$ and $F_{\text{After}}$ as a transition density plot (TDP). **(3)** The duration that a particular FRET state exists in $F_{\text{Before}}$ before transition to $F_{\text{After}}$ is defined as the residence time. The residence time for each transition class described in (1) can be collated and presented as a bar plot showing the mean ± standard error of the mean. T-1, T-2 etc. refers to different experimental treatments.

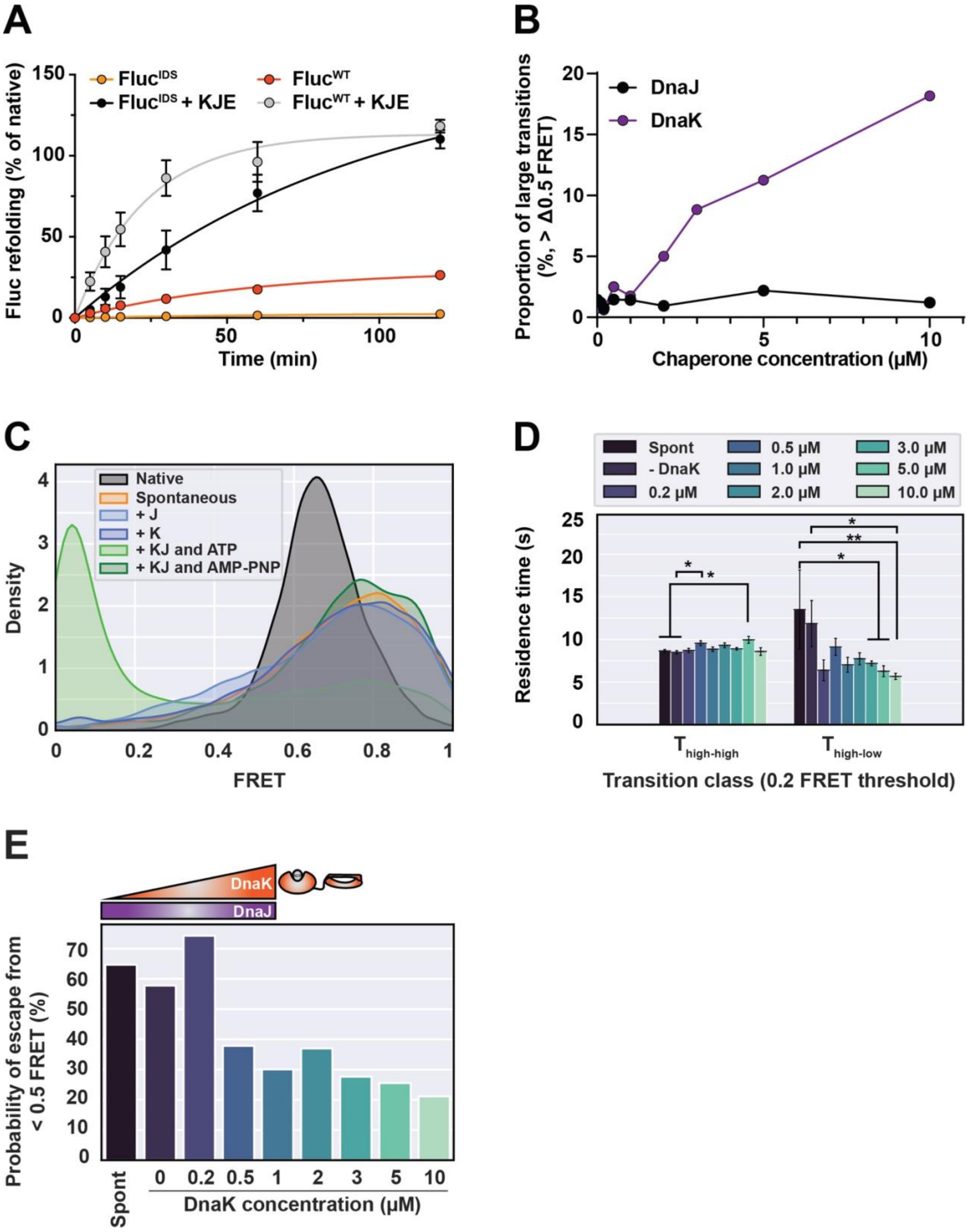


**Supplementary Figure 3: High concentrations of DnaK promote significant conformational rearrangement of Fluc$^{IDS}$ and prevents collapse of the client. (A)** Fluc$^{IDS}$ or Fluc$^{WT}$ in 5 M GdHCl was diluted 100-fold (10 nM final concentration) into refolding buffer alone (i.e., spontaneous, *Fluc$^{IDS}$* or *Fluc$^{WT}$*) or buffer supplemented with molecular chaperones. Chaperone-assisted refolding was performed with DnaK (3 µM), DnaJ (1 µM), GrpE (1.5 µM) and ATP (5 mM) (+ *KJE*). Refolding was performed at 25˚C and aliquots were taken at various timepoints for detection of luminescence. All luminescence values were normalized to that of native (non-denatured) Fluc$^{IDS}$ or Fluc$^{WT}$ samples. Data shown is the mean ± standard error of the mean from three independent experiments. **(B)** Misfolded $Fluc^{IDS}$ was incubated with increasing concentrations of DnaJ (0 - 10 μM, *DnaJ*) or in the presence of DnaJ (0.2 μM), ATP (5 mM) and increasing concentrations of DnaK (0 – 10 μM, *DnaK*) and the FRET efficiency measured. The proportion of transitions that were large (> Δ0.5 FRET in magnitude) was then calculated and plotted. **(C)** FRET efficiency histograms of misfolded $Fluc^{IDS}$ incubated in the presence of either DnaJ (0.2 μM) or DnaK (3 μM) alone or in the presence of DnaJ (0.2 μM), DnaK (3 μM) and either ATP (5 mM) or AMP-PNP (5 mM). **(D)** Bar plots showing the mean residence time ± standard error of the mean for $T_{high\text{-}high}$ and $T_{high\text{-}low}$ transitions, where a FRET threshold of 0.2 is set to determine 'high' or 'low' states. A two-way ANOVA statistical analysis with Tukey's multiple-comparisons post-hoc test was performed to determine statistically significant differences in residence times between treatment groups within each transition class. * and ** indicates statistical significance with $p \leq 0.05$ and 0.01, respectively. The absence of markers indicates no significant difference ($p > 0.05$). **(E)** The probability that a $Fluc^{IDS}$ molecule in the presence of DnaJ (0.2 μM) and increasing concentrations of DnaK (0 – 10 μM) will transition to a high-FRET state (> 0.5) from a low-FRET state (< 0.5).

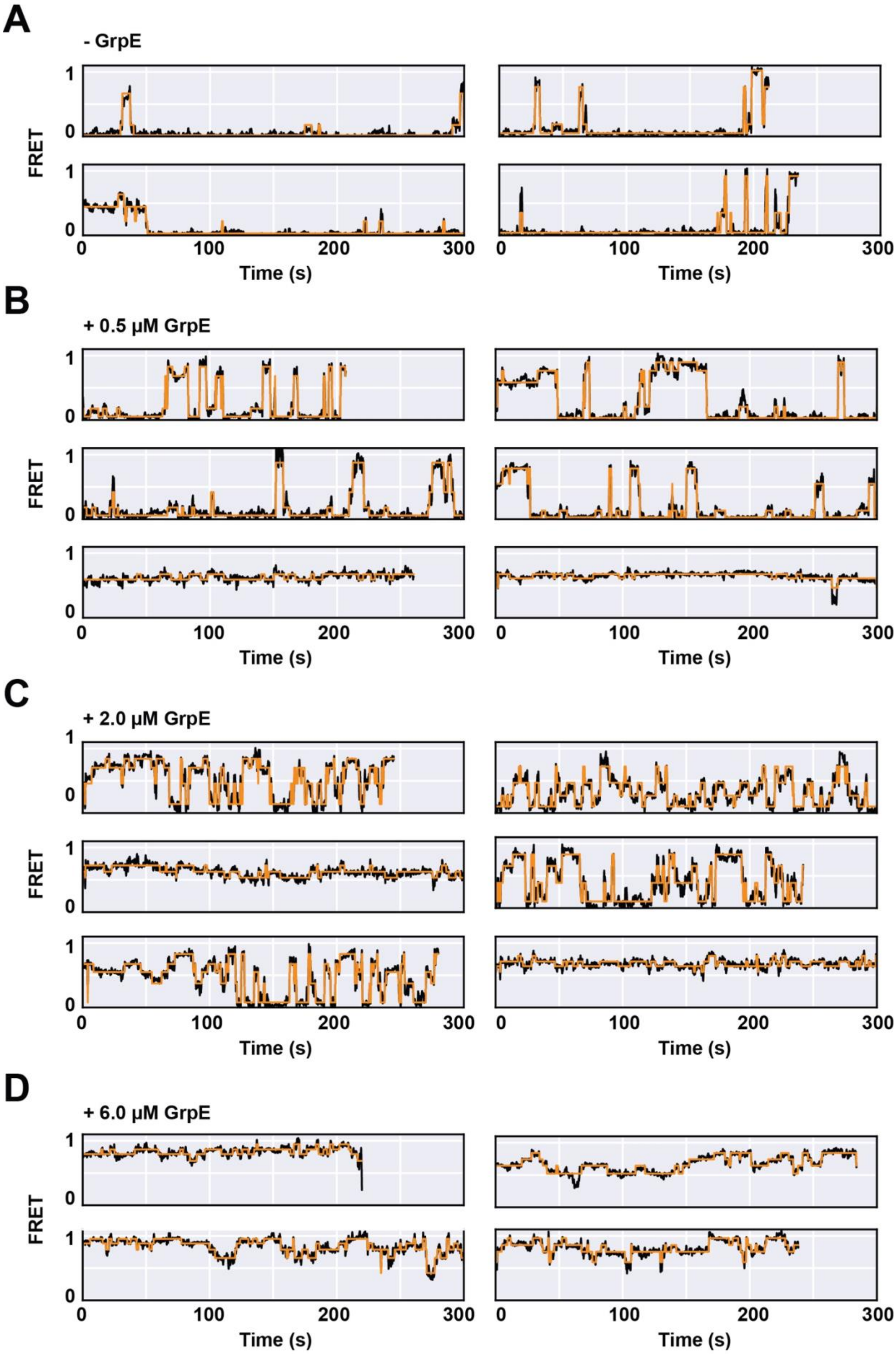
A
- GrpE
B
+ 0.5 µM GrpE
C
+ 2.0 µM GrpE
D
+ 6.0 µM GrpE
FRET
Time (s)
0
1
100
200
300

**Supplementary Figure 4: Example smFRET trajectories of individual Fluc$^{IDS}$ molecules during chaperone-assisted folding.** AF555/647-labelled Fluc$^{IDS}$ was immobilized to a coverslip surface and incubated with DnaK (9 µM), DnaJ (0.6 µM), ATP (5 mM) in the absence or presence of increasing concentrations of GrpE (0.5 – 6 µM). The FRET efficiency of individual Fluc$^{IDS}$ molecules was measured for 36 min following the addition of GrpE. FRET intensity traces of Fluc$^{IDS}$ incubated in **(A)** the absence of GrpE or in the presence of **(B)** 0.5 µM GrpE, **(C)** 2.0 µM GrpE or **(D)** 6.0 µM GrpE.

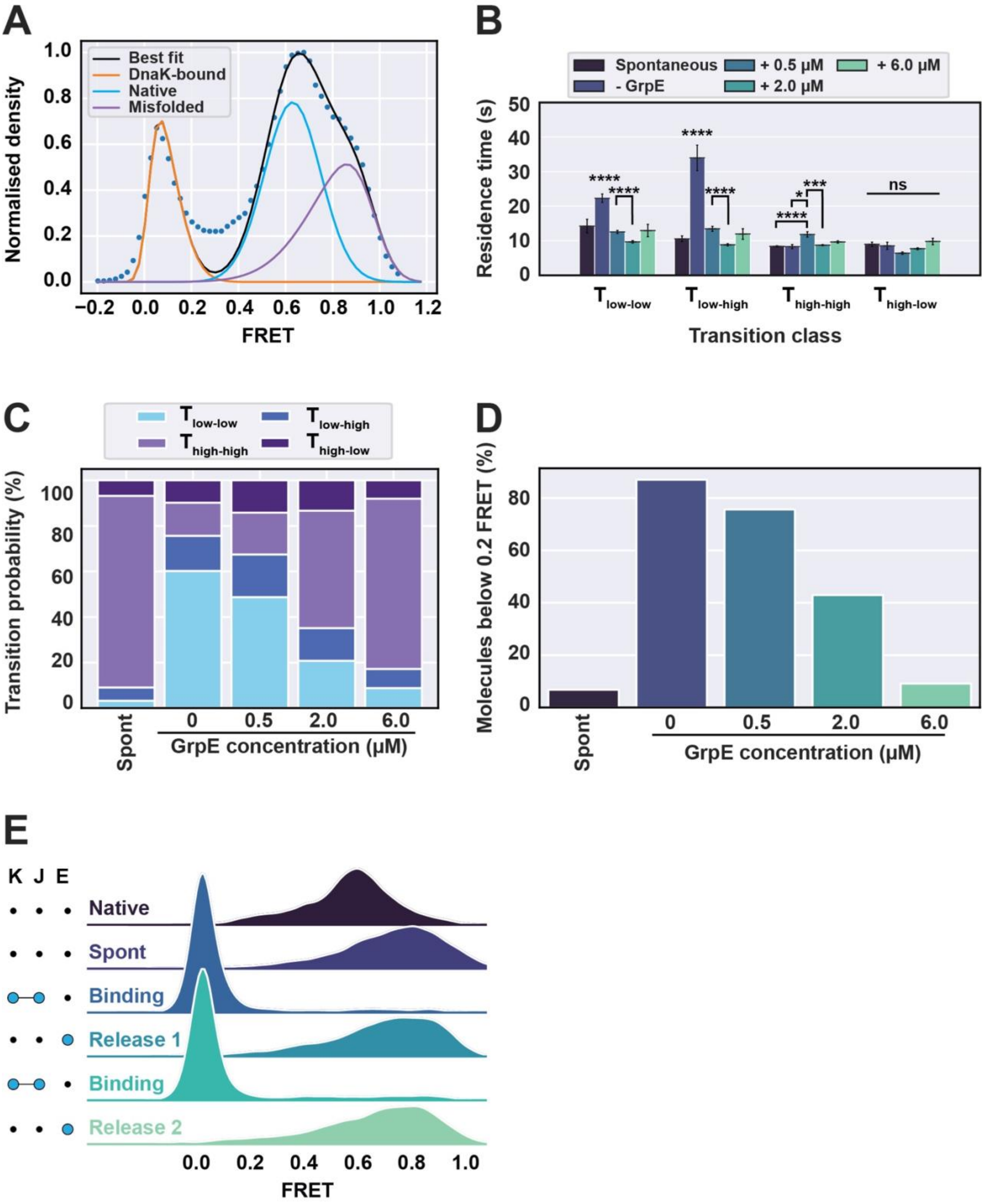

**Supplementary Figure 5: GrpE promotes protein folding by regulating the lifetime of the DnaK-Fluc$^{IDS}$ complex and facilitating multiple chaperone binding-and-release cycles.** Fluc$^{IDS}$ was incubated in the presence of DnaJ (0.6 µM), DnaK (9 µM) and ATP (5 mM) supplemented with increasing concentrations of GrpE (0 – 6 µM). FRET efficiency data was measured for 36 min following the addition of GrpE. **(A)** An example FRET efficiency histogram dataset with the multiple gaussian model fits used to determine the proportion of Fluc$^{IDS}$ states when incubated in the presence of molecular chaperones. **(B)** Bar plots showing the mean residence time ± standard error of the mean for each transition class. A two-way ANOVA statistical analysis with Tukey's multiple-comparisons post-hoc test was performed to determine statistically significant differences in residence times between treatment groups within each transition class. **(C)** The transition probability of each transition class. **(D)** Bar plot of the percentage of molecules that visit the DnaK-bound state, defined as < 0.2 FRET. **(E)** To determine if a single round of chaperone binding-and-release is sufficient for refolding, misfolded Fluc$^{IDS}$ (*spont*) was alternatively incubated in the presence of DnaJ (1 µM), DnaK (15 µM) and ATP (5 mM) (*binding*) or GrpE (10 µM) alone (*release*) and repeated once. The FRET efficiency was measured from individual Fluc$^{IDS}$ molecules for each condition for 30 min and presented as a FRET efficiency histogram. When statistical tests were performed, *, **, *** and **** indicates statistical significance with $p \leq 0.05$, 0.01, 0.001 and 0.0001, respectively. ns or the absence of markers indicates no significant difference ($p > 0.05$).
